# Identification of *cis* elements for spatio-temporal control of DNA replication

**DOI:** 10.1101/285650

**Authors:** Jiao Sima, Abhijit Chakraborty, Vishnu Dileep, Marco Michalski, Juan Carlos Rivera-Mulia, Claudia Trevilla-Garcia, Kyle N. Klein, Daniel Bartlett, Brian K. Washburn, Michelle T. Paulsen, Daniel Vera, Elphège P. Nora, Katerina Kraft, Stefan Mundlos, Benoit G. Bruneau, Mats Ljungman, Peter Fraser, Ferhat Ay, David M. Gilbert

## Abstract

The temporal order of DNA replication (replication timing, RT) is highly coupled with genome architecture, but *cis*-elements regulating spatio-temporal control of replication have remained elusive. We performed an extensive series of CRISPR mediated deletions and inversions and high-resolution capture Hi-C of a pluripotency associated domain (DppA2/4) in mouse embryonic stem cells. Whereas CTCF mediated loops and chromatin domain boundaries were dispensable, deletion of three intra-domain prominent CTCF-independent 3D contact sites caused a domain-wide delay in RT, shift in sub-nuclear chromatin compartment and loss of transcriptional activity, These “early replication control elements” (ERCEs) display prominent chromatin features resembling enhancers/promoters and individual and pair-wise deletions of the ERCEs confirmed their partial redundancy and interdependency in controlling domain-wide RT and transcription. Our results demonstrate that discrete *cis*-regulatory elements mediate domain-wide RT, chromatin compartmentalization, and transcription, representing a major advance in dissecting the relationship between genome structure and function.

**Highlights:**

- *cis*-elements (ERCEs) regulate large scale chromosome structure and function
- Multiple ERCEs cooperatively control domain-wide replication
- ERCEs harbor prominent active chromatin features and form CTCF-independent loops
- ERCEs enable genetic dissection of large-scale chromosome structure-function.

## INTRODUCTION

Eukaryotic DNA replication follows a defined temporal order (replication timing; RT) that, in mammals, is mediated by the coordinated firing of clusters of origins within 400-800kb chromosomal units (replication domains; RDs) at different times during S phase (Rivera-Mulia and Gilbert, 2016a). RT is developmentally regulated and correlated with transcriptional potential (Rivera-Mulia and Gilbert, 2016b), and distinct mutational signatures (Sima and Gilbert, 2014). Defects in RT have been reported in several diseases, correlated with mis-regulation of genes and associated with defects in chromosome condensation, sister chromatid cohesion, and genome instability (Koren et al., 2014; Platt et al., 2018; Sasaki et al., 2017). RT is also closely correlated to large-scale genome architectural organization within the nucleus. Early cytological studies demonstrated that sites of briefly labeled DNA synthesis could be visualized as punctate foci that behaved as stable units of chromosome structure throughout multiple cell cycles (Solovei et al., 2016). These units were found to be spatio-temporally segregated in the nucleus such that early replicating foci are dispersed throughout the interior of the nucleus, while late replicating foci are in close proximity to the nuclear lamina and nucleoli.

The structure of RDs and their spatio-temporal segregation can now be mapped molecularly. High throughput chromosome conformation capture (Hi-C) reveals as a first principle component two major compartments of chromatin interactions termed A (open) and B (closed) (Dixon et al., 2012; Lieberman-Aiden et al., 2009) that strongly correlate with genome-wide molecular maps of RT (Ryba et al., 2010; Yaffe et al., 2010), as well as proximity to the nuclear lamina and nucleoli (Pope et al., 2014). Hi-C also identifies units of self-interacting chromatin called topologically associating domains or TADs (Dixon et al., 2012; Nora et al., 2012) that were later shown to correspond to RDs (Pope et al., 2014). Selected groups of RD/TADs coordinately change their RT, chromatin compartment and sub-nuclear position during differentiation (Dixon et al., 2015; Ryba et al., 2010; Takebayashi et al., 2012) and both RD/TAD structure and A/B compartment organization are dismantled during mitosis and re-assembled during early G1 phase, coincident with the re-establishment of a distinct RT program (Dileep et al., 2015a). RD/TADs are also units of gene regulation, histone modifications and genome stability (Schmitt et al., 2016).

The molecular mechanisms regulating the spatial and temporal segregation of chromatin are poorly understood. Perturbations of many chromatin regulatory proteins have only partial or localized effects on RT or A/B Hi-C compartments (Dileep et al., 2015b; Foti et al., 2016; Nora et al., 2017; Rao et al., 2017; Weintraub et al., 2017) and *cis*-acting elements regulating mammalian RT or A/B compartments have not been identified to date and it has not been clear whether a role for DNA sequence in such large-scale regulation would be via discrete elements or more complex features such as overall nucleotide composition or secondary structure. Early studies suggested that the β-globin Locus Control Region (LCR) is sufficient to regulate RT at ectopic loci (Goren et al., 2008; Hassan-Zadeh et al., 2012; Simon et al., 2001), but targeted deletion of the LCR at its native locus had no effect on RT (Cimbora et al., 2000). Other studies have identified DNA segments that regulate sub-nuclear localization (Hassan-Zadeh et al., 2012; van de Werken et al., 2017; Zullo et al., 2012), and a human chromosome has been shown to retain its RT regulation when carried in a mouse host (Pope et al., 2012), consistent with a role for the underlying DNA sequence in spatio-temporal regulation. To search for putative *cis*-acting elements, we performed an extensive series of CRISPR-mediated (clustered regularly interspaced short palindromic repeats) deletions and inversions at a pluripotency-associated RD/TAD in mESCs. Our results revealed three partially redundant, inter-dependent and strongly interacting (capture Hi-C) *cis*-regulatory elements responsible for early replication and chromatin compartmentalization. These elements have properties of super-enhancers, and play roles in transcription within this domain, but are independent of the replication origins themselves. Together our results demonstrate the existence of *cis*-acting elements of large-scale spatio-temporal control of genome structure and function and provide a means to identify such elements at other domains genome-wide and investigate their roles in large-scale chromosome architecture and function.

## RESULTS

### Epigenetic landscape of the DppA2/4 replication domain

To identify *cis*-acting elements of replication domain structure, we employed CRISPR/Cas9 technology to generate a series of deletions and inversions at a developmentally regulated replication domain (heretofore termed the DppA2/4 domain) located on chr16 in mouse ESCs (**Figure 1a and Supplemental Figure 1**). This domain contains 3 active genes: DppA4, DppA2 and Morc1/Morc (**Figure 1a**). DppA4 and DppA2 are oncogenes (Tung et al., 2013) and important ESC markers, and KO mice are embryonic lethal (Du et al., 2010; Ivanova et al., 2006; Madan et al., 2009; Masaki et al., 2007; Nakamura et al., 2011). Morc1 is a multi-functional gene that is key for male meiosis and spermatogenesis, and is involved in bone metabolism, cancer development (Hong et al., 2017) and major depressive disorder (Schmidt et al., 2016). It is highly expressed in ESCs and interestingly, contains an alternative transcription start site (TSS) 180kb downstream (Morc) of the original TSS detected by Cap Analysis of Gene Expression (CAGE) (Arner et al., 2015) (**Supplemental Figure 1b**). This domain is one of a select group of domains that switch from early to late during loss of pluripotency and remain late replicating in all subsequent developmental stages (Hiratani et al., 2010) (**Supplemental Figure 1b**). Genes within this class of domains are particularly difficult to reprogram back to the early replicating / transcriptionally active state when making induced pluripotent stem cells (iPSCs) (Hiratani et al., 2010). Early to late RT regulation of the DppA2/4 domain is conserved in humans (Dileep et al., 2015a), and coincides with repression of all three genes, spatial compaction of the domain, movement to the nuclear periphery and a change in Hi-C compartment consistent with increased interactions to neighboring late replicating domains (Hiratani et al., 2010; Takebayashi et al., 2012). The DppA2/4 domain is maintained as an inter-LAD in ESCs as defined by laminB1 DamID and is flanked on both sides by ~3Mb LADs, the left of which is a constitutively late replicating gene desert (**Supplemental Figure 1b**). The DppA2/4 domain aligns with a TAD whose boundaries are occupied by CTCF and cohesin proteins (**Supplemental Figure 1b**), and contains putative internal regulatory elements with active histone marks, making it an excellent candidate to address TAD/RD structure and function.

**Figure 1.**
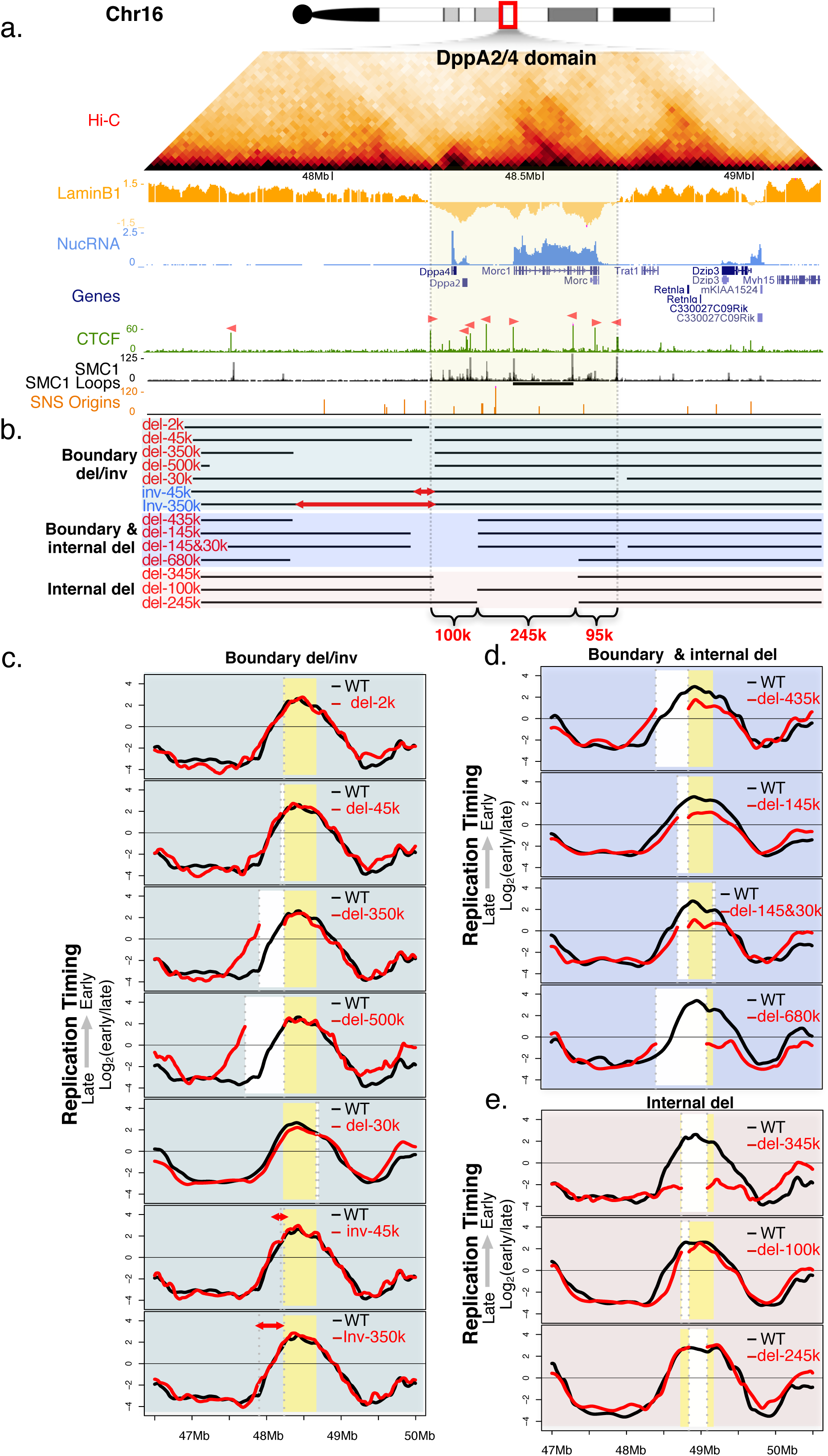
Internal segments contribute partially to early replication. a. 3D structure and chromatin features of the DppA2/4 domain. Presented in order are Hi-C heat map, LaminB1 DamID (LaminB1), nuclear RNA (NucRNA), reference genes (Genes), CCCTC-binding factor ChIP (CTCF), SMC1 ChIP signal in ChIAPET datasets (SMC1), SMC1 ChIAPET identified loops (SMC1 loops, horizontal bar in black), and short nascent strand mapped replication origins (SNS origins) in mESCs. CTCF binding site orientation is indicated as pink triangles above the CTCF ChIP track (See Supplementary Methods). The grey vertical lines indicate the domain boundaries. b. Diagram indicating the positions of the boundary deletions or inversions, boundary and internal deletions, and internal deletions. c,d,e. RT profiles of corresponding deletions or inversions (red lines) as compared to WT control (black lines). The DppA2/4 replication domain is highlighted in yellow; deleted regions are masked in white; inversions are indicated by red arrows; and breakpoints are indicated by grey dashed lines.

### CTCF-associated TAD boundaries and CTCF protein are dispensable for maintaining RT of the DppA2/4 domain

We first investigated the role of CTCF in regulating RT and TAD integrity by deleting or inverting sequences harboring the CTCF binding sites at the boundaries (**Figure 1b & Supplemental Figure 1**). The DppA2/4 domain is flanked by CTCF peaks, with the right border peak co-localizing with cohesin. There is also a CTCF/cohesin-mediated loop within the domain revealed by Smc1 CHIA-PET analysis (Dowen et al., 2014) (**Figure 1a**). Deletions of 2kb, 45kb, 335kb, or 500kb, encompassing the CTCF associated left boundary, or 30kb of the CTCF/cohesin co-occupied right boundary had no effect on RT of the remaining portion of the domain (**Figure 1c**). Next, we tested whether inversion of the orientation-dependent CTCF sites could disrupt contact domain structure by creating new loops in the reverse orientation as observed before (Lupiáñez et al., 2015). However, inversion of 45kb and 335kb harboring the CTCF associated left boundary also had no effect on RT (**Figure 1c**). Circularized chromosome conformation capture (4C) analysis using a bait within the domain following the boundary deletions did not detect evidence of change in preferential chromatin interactions (**Supplementary Figure 2**). These results suggest that neither TAD structure nor RT require CTCF-associated TAD boundaries at the DppA2/4 domain.

To assess the role of CTCF on RT genome wide, we profiled RT in a recently developed cell line in which CTCF protein levels are acutely depleted using the auxin-inducible degron (AID) system (Nora et al., 2017) (**Supplementary Figure 3**). CTCF depletion resulted in very few minor changes in RT (0.44% of the genome with RT delay; 0.14% with RT advance) that preferentially occurred at domains with developmentally regulated RT (**Supplementary Figure 3**). This is consistent with the finding that CTCF depletion also had no impact on genome compartments (Nora et al., 2017). In fact, the DppA2/4 domain boundaries are still present (**Supplementary Figure 3e**), and DppA2/4 RT was not affected in cells depleted of CTCF (Nora et al., 2017) (**Supplementary Figure 3d**). Taken together, these results demonstrate that CTCF plays very little role in DppA2/4 domain boundaries, RT or in the global RT program.

### Multiple internal DNA sequences contribute partially to early replication of the DppA2/4 domain

Since CTCF associated boundary deletions or inversions had no effect on RT, we hypothesized that elements internal to the domain may contribute to early replication. To avoid potential loss of pluripotency caused by homozygous deletion of DppA2/4 genes and to allow a direct comparison between the deleted and wild-type allele within the same datasets, we generated heterozygous deletions in mESCs derived from a cross between *M. casteneus* (CAST/Ei) and *M. musculus* (129/sv) (Dupont et al., 2016), from which reads can be distinguished based on single nucleotide polymorphisms (SNPs) at an average density of one per 150 base pairs. Some deletions were also performed in mESC line V6.5, derived from a cross between the *129/sv* and *C57BL/6* strains of *M. musculus* and harboring SNPs at an average density of one per kilobase (kb). A detailed list of which deletions were made in each of the cell lines studied in this report is catalogued in **Supplementary Table 1**.

We first extended two prior boundary deletions (335k, 45k) by an additional 100kb into the interior of the domain yielding 435kb and 145kb in total (**Figure 1b**). As shown in **Figure 1d**, this extra 100kb deletion including the DppA2 and DppA4 genes within the domain caused a slight delay in RT. Deletion of 30kb encompassing the right boundary in the context of the 145kb did not cause any further delay, consistent with no role for the boundary in RT. We next further extended these deletions 245 kb into the DppA2/4 domain (680 kb deletion) removing the Morc1 transcriptional start site (TSS), additional sites of active histone marks, an internal CTCF/cohesin loop anchor, most of the inter-LAD chromatin as well as a cluster of replication origins (Cayrou et al., 2015). This invasive deletion caused a substantial delay in RT of the remaining 95kb of the Dppa2/4 domain, which remained nonetheless an early replicating peak. Deleting only the 345kb interior portion of the 680kb invasive deletion, which left the CTCF boundaries intact, had a similar effect on RT as the 680kb deletion, confirming that the control elements responsible for this RT shift are interior to the domain (**Figure 1e**). Attempts to dissect the 345kb segment by making 100kb and 245kb deletions, as well as an extensive series of smaller deletions (**Figure 1e** and **Supplementary Figure 1**), had little to no detectable effect on DppA2/4 domain RT. Altogether, this series of deletions suggests the existence of multiple redundant or cooperative segments within the interior of the DppA2/4 domain that are necessary for early replication of DppA2/4 domain.

### Large CRISPR inversions demonstrate sufficiency of internal segments for early replication

To determine whether the segments identified above are sufficient to dictate early replication in novel structural configurations, we generated a 680kb inversion containing 345kb of the DppA2/4 domain and 335kb of the adjacent late replicating gene desert, separating the 345kb DppA2/4 domain sequences from the remaining 95kb inter-LAD and the original right boundary (**Figure 2a**). Strikingly, the 345kb DppA2/4 segment was sufficient to recapitulate a nearly perfectly inverted early replication pattern relative to the WT allele, and the remaining 95kb persisted as the locally earliest replicating segment, replicating in mid-S phase (**Figure 2b**). Similarly, a 435K inversion containing 100k of the DppA2/4 sequences was able to maintain its early RT (**Figure 2c**). These results suggest the existence of *cis*-regulatory elements that are both necessary and sufficient to maintain early RT independent of their native boundary. To confirm these findings at an independent locus, we analyzed replication timing in a series of previously published deletions and inversions at a constitutively early replicating domain (Wnt6/Ihh) on chr1 in mESCs (Lupiáñez et al., 2015) (**Supplementary Figure 4**). Early and late replication of both the Wnt6/Ihh and Epha4 domains was largely retained after removing large segments containing the boundary in Dbf/+ cell line (**Supplementary Figure 4b**). In the case of the 1.07Mb inversion InvF/InvF, the Ihh domain sequences remained early replicating, consistent with the findings in the DppA2/4 domain. These results provide further support for the existence of *cis*-regulatory elements of early replication that are independent of TAD boundaries and demonstrate that they are not restricted to the DppA2/4 locus or to developmental domains.

**Figure 2.**
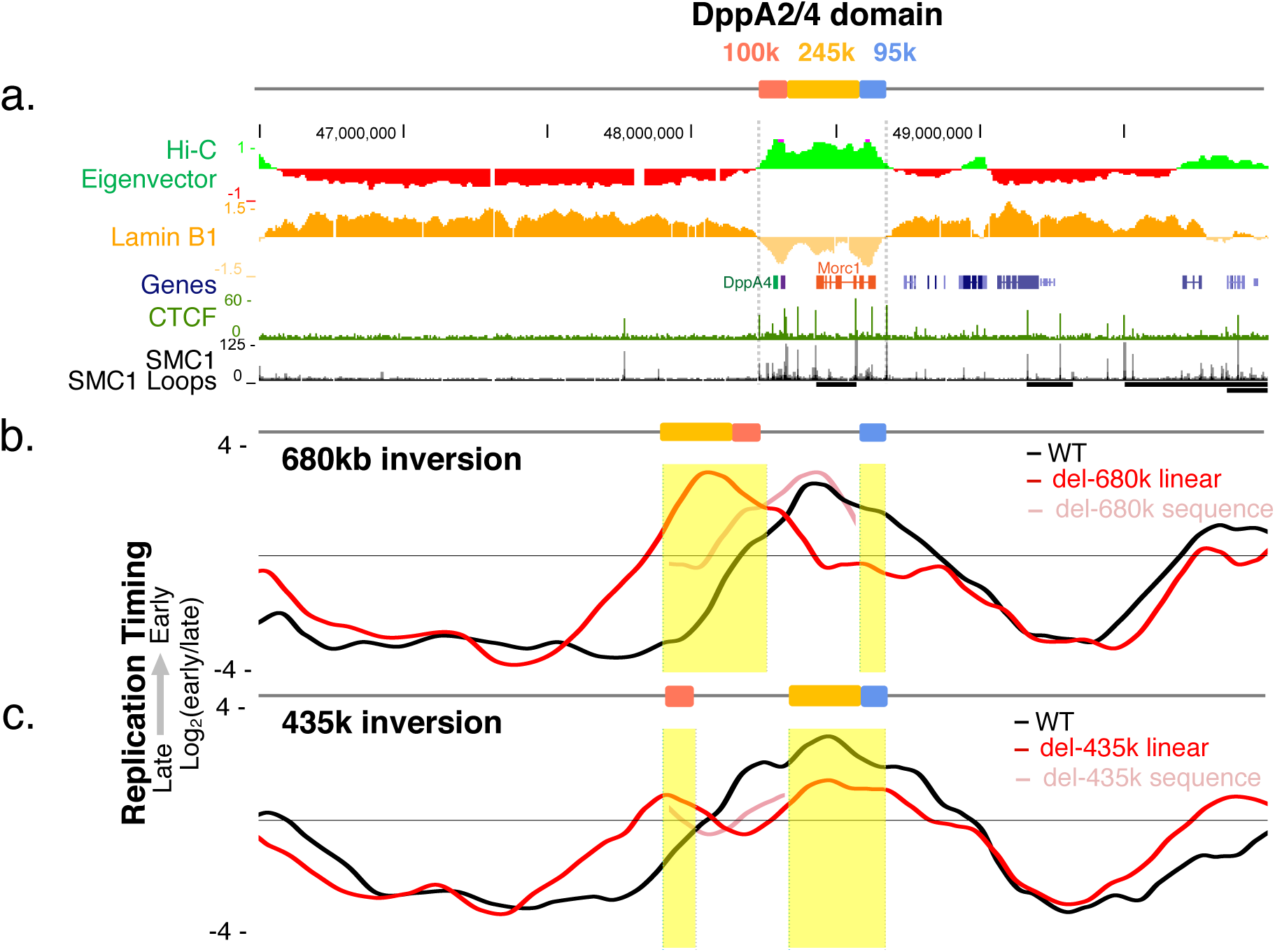
Large-scale inversions demonstrate sufficiency of internal segments for early replication. a. Schematic presentation of genomic features near DppA2/4 domain. b,c. RT profile for the 680kb (b) and 435kb (d) inversions with WT allele plotted in black, inverted allele in red in it’s actual linear distance, and the inverted allele in pink as it would appear with WT coordinates.

### Identification of Early Replication Control Elements (ERCEs)

As the redundancy among different segments impeded our ability to define *cis*-elements in RT regulation using a systematic series of deletions, we turned to a candidate approach to identify the *cis*-acting elements. Since RT strongly correlates with chromatin 3D compartmentalization (Dileep et al., 2015a; Pope et al., 2014; Ryba et al., 2010), we reasoned that a high-resolution 3D contact map of the DppA2/4 domain could potentially reveal features of domain structure consistent with the effects of *cis*-deletions and inversions described above. We designed a set of capture Hi-C (cHi-C) (Dryden et al., 2014) oligonucleotides hybridizing to ~8,000 SNP-containing MboI fragments covering ~5Mb region surrounding DppA2/4 domain (**Figure 3a**). The library was sequenced with ~3 million valid read pairs per CAST/Ei vs. 129/sv allele, equivalent to 1.6 billion read pairs per genome for a genome-wide approach (e.g., Hi-C), providing high resolution contact maps of each homologue. The resulting contact maps at 5kb resolution recapitulated the dramatic local folding pattern shift during differentiation from ESCs to NPCs (Takebayashi et al., 2012) (**Supplementary Figure 5a&b**). To identify genomic sites that exhibit statistically significant contacts within the DppA2/4 domain we applied Fit-Hi-C (Ay et al., 2014), which jointly models the distance decay and other technical biases in Hi-C datasets (GC content, fragment length, mappability, and capture efficiency). Several sites, including the pair previously identified by cohesin ChIA-PET (Dowen et al., 2014), emerged as the most prominent long-range contacts within the domain (FDR of 0.1%) (**Figure 3c and Supplementary Figure 5**). As CTCF boundary deletion and CTCF depletion had no effect on DppA2/4 RT (**Figure 1c & Supplementary Figure 3**), we focused on the 3 sites that form CTCF-independent loops. These sites were also enriched in multiple active epigenetic features (DNase1 HS, P300, H3K27ac, H3K4m1, H3K4m3 or Med1, a TSS and master regulator transcription factor binding peaks) that are hallmarks for enhancers, and associated with early replication (**Figure 3b & Supplementary Figure 6**). A site near the DppA2 gene also corresponds to the P300 peak of an ESC-annotated super-enhancer (Dowen et al., 2014; Khan and Zhang, 2016), virtually all (229/231) of which we found to be early replicating. These 3 sites, which we named a, b, and c in their linear order on the chromosome, are distributed such that each of the 100k, 245k and 95k segments from our previous deletion series contains one of them (**Figure 3b**). Strikingly, targeted deletion of all 3 sites caused a complete switch from early to late replication of the Dppa2/4 domain, similar in magnitude to the RT shift seen during NPC differentiation (**Figure 3d**). This phenotype was reproduced in 2 independent CRPISR deletion clones, validating that we have depleted all essential elements contributing to early replication of the DppA2/4 domain. Given the discrete nature of these elements, we have designated them as Early Replication Control Elements (ERCEs). Notably, two sites outside of ERCEs (X and Y in **Figure 3b**) that either lack a full profile of chromatin marks or do not interact strongly with ERCEs were neither necessary nor sufficient for early replication. Site X contains the TSS of the highly transcribed gene DppA4, forms a weaker loop with site a, but lacks some of the chromatin features of ERCEs. Site Y contains similar chromatin features as ERCEs but does not form prominent loops and does not contain a TSS. These two sites suggest the importance of both a specific chromatin composition and the ability to form strong CTCF-independent interactions for ERCE activity. In addition, these segments do not overlap with the majority of efficient replication initiation sites (Cayrou et al., 2015) (SNS origin track in **Figure 3b**), consistent with the separation of temporal control from the sites of initiation (Dileep et al., 2015b).

**Figure 3.**
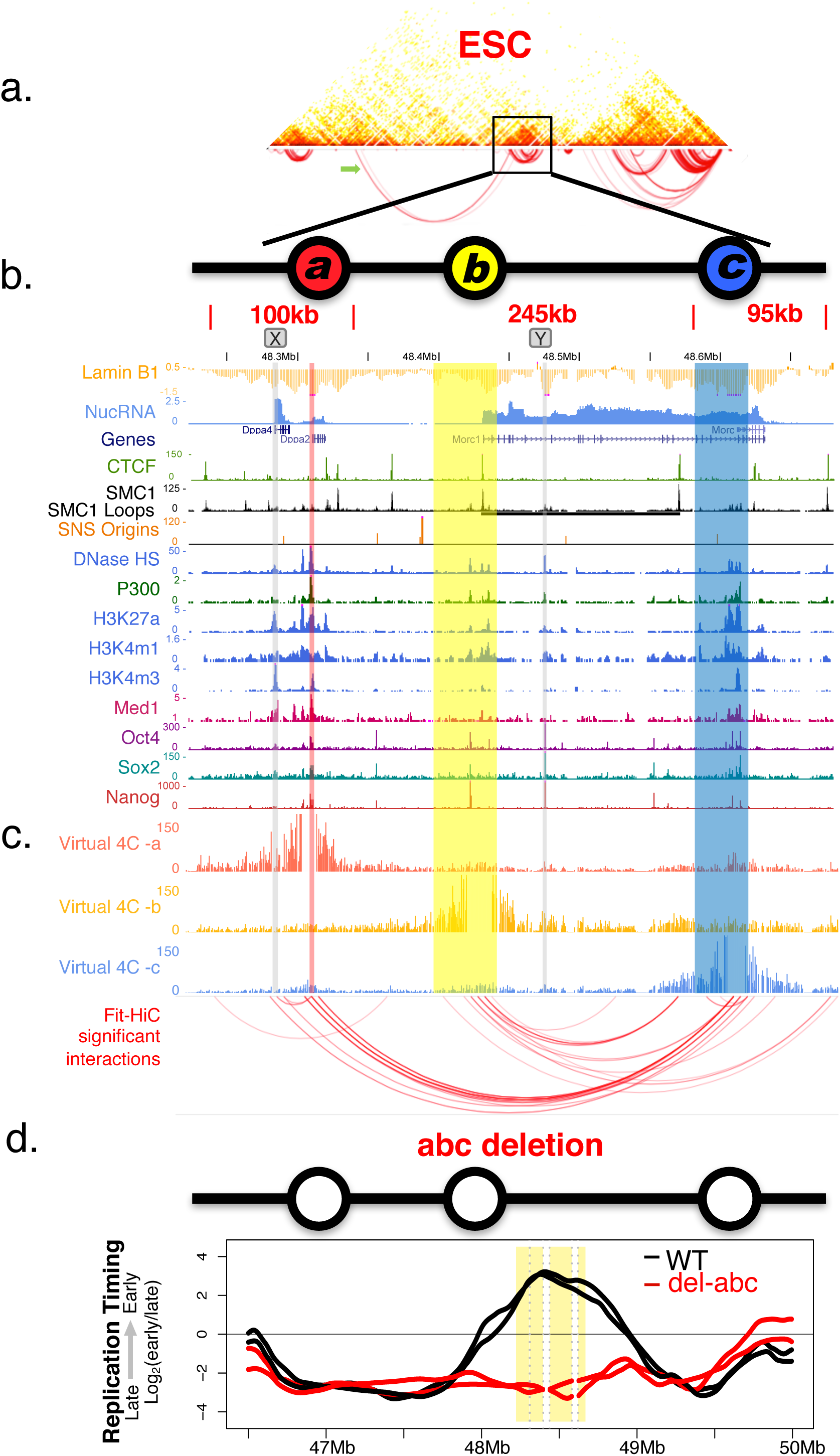
Identification of Early Replication Control Elements (ERCEs). a. A high-resolution capture Hi-C heatmap of the ~5Mb region including DppA2/4 domain with underlying arc plot of the Fit-HiC significant interaction pairs. The green arrow indicates the long-range interaction described in Figure 4a. b. Chromatin features of the DppA2/4 domain highlighting three sites (a,b,c) enriched in various active marks. Site X and Y are examples other than ERCEs that are neither necessary nor sufficient for the domain early replication. c. Site a,b,c emerge as the strongest CTCF/cohesin-independent interactions within the domain. Virtual 4C profiles from the viewpoints of either site a, b or c are plotted for DppA2/4 domain at the resolution of MboI restriction fragments. Bait regions are removed from the plots. Significant interactions identified by Fit-HiC are presented as red arcs (FDR >0.01). d. RT profile for abc triple deletion showing a complete shift to late replication of the DppA2/4 domain, with mutant allele plotted in red, and WT allele plotted in black. Profiles from two different clones are presented.

### ERCEs also control A/B compartmentalization

ERCEs not only form the most robust intra-domain interactions within the domain, but also they interact strongly with ERCE–like elements outside the domain. For example, site a forms a strong long-range interaction with the nectin3/pvrl3 promoter (**Figure 3a and 4a**), which is also a locally earliest replicating segment, resides in compartment A and is deposited with similar chromatin composition as site a, b or c (**Figure 4b**). In addition, the interactions between ERCEs are CTCF independent as they persist as strong interaction sites after CTCF protein depletion (**Supplementary Figure 5d**). These CTCF-independent intra- and inter-domain interactions suggest a potential role for ERCEs in driving A/B compartmentalization. To determine the effect of ERCE deletion on compartmentalization, we performed 4C-seq with a bait positioned between site b and site c present in both WT and deleted alleles. Allele specific 4C-seq results revealed that the abc deleted allele switched to interact with late replicating chromatin in compartment B, to a similar degree as in NPCs (**Figure 4c&d**). Therefore, ERCEs are responsible for both early replication and sub-nuclear compartmentalization.

**Figure 4.**
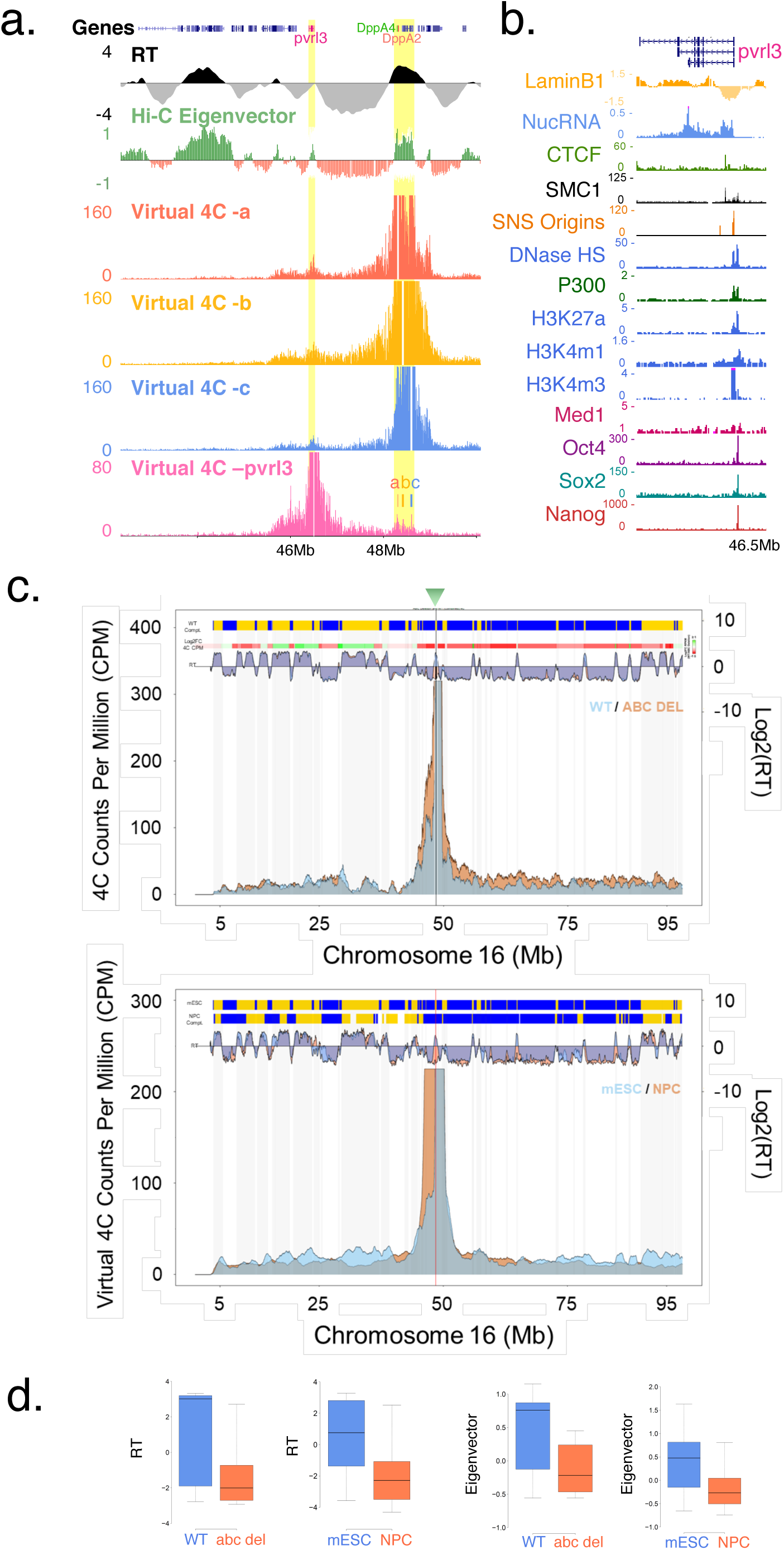
ERCE mediated interactions may drive A/B compartmentalization. a. ERCEs interact with ERCE-like elements outside DppA2/4 domain. Virtual 4C profiles generated from Capture Hi-C data (chr16:45488946-50879849 is enriched) are plotted from the viewpoints of either site a, b, c or the ERCE-like element pvlr3/nectin3 promoter region upstream of DppA2/4 domain. Bait regions are removed form the plots. The pvlr3/nectin3 promoter region is locally the earliest replicating point and associated with compartment A. The pvlr3/nectin3 promoter region and DppA2/4 domain are shaded in yellow. b. The pvlr3/nectin3 promoter region is decorated with ERCE-like chromatin features. c. Plots of smoothed 4C contact counts from a viewpoint (green triangle) inside DppA2/4 domain on the abc deleted (orange) and WT allele (blue). Hi-C eigenvector, log2 ratio of 4C fold changes (WT/abc del), and RT are plotted above the 4C profile. Similarly, virtual 4C contact counts from the same viewpoint are plotted for ESC (blue) and NPC (orange). Increased interactions to compartment B/late replicating regions can be observed in both abc deleted alleles and NPC. c. Boxplots quantifying RT or Hi-C eigenvector distribution of 4C significant contacts in abc deletion or NPC differentiation.

### Interdependence of ERCEs in spatio-temporal control

We next performed individual and pair-wise deletions of sites a, b and c to further investigate the cooperative effect of these sites (**Figure 5**). Single deletion of either site a or b did not significantly alter RT, consistent with results of the larger deletions containing either a or b in the previous series (100k, 28.5k and 28k for site a, 245k for site b; **Figure 1e** and **Supplementary Figure 1**). However, when site c alone was deleted, the right section of the domain acquired the sloping pattern of an initiation-free timing transition region (TTR) (Dileep et al., 2015b) with a slope of progressively later RT, while the left section containing a and b remained early. These results imply that the domain is divided into two segments in which early RT of the right requires site c, while the left section is controlled by a or b. Interestingly, we found that all pair-wise deletions gave rise to a similar pattern with the ERCE remaining in the domain being the locally earliest replicating segment. For example, the a-b double deletion led to late replication of the left side of the domain while the right portion encompassing site c continued to replicate in mid-S phase. This result was similar to the previous large deletion encompassing site a and b (680 kb and 345kb deletion) (**Figure 1d-e**), indicating that a and b can account for the effects of those larger deletions. Together, this set of deletions reveals the partial redundancy and inter-dependence between the three major sites in giving rise to domain-wide early replication of the DppA2/4 domain, accounting for the complexity in the results of the unbiased nested deletion approach (**Figure 1** and **Supplementary Figure 1**).

**Figure 5.**
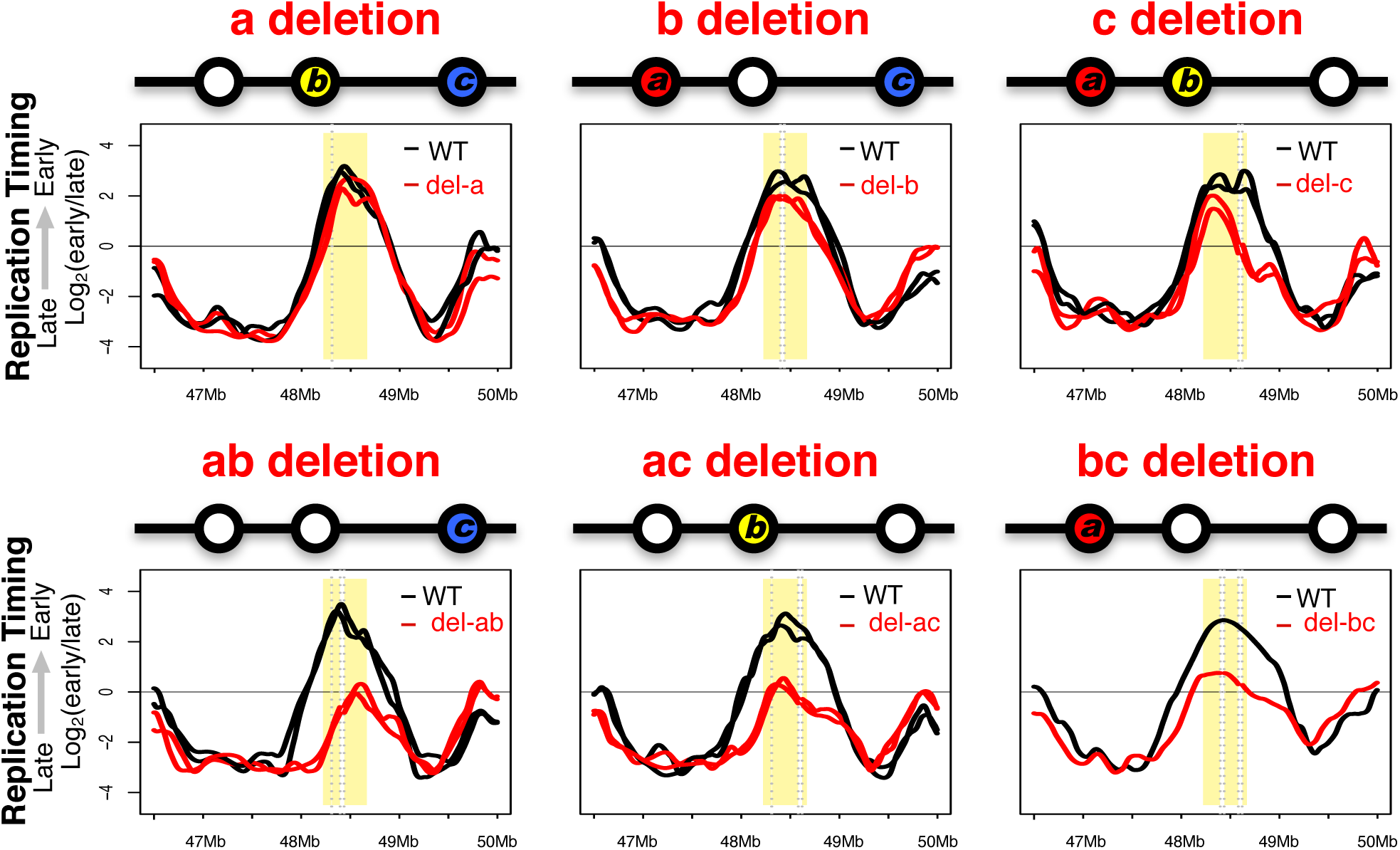
Individual or pair-wise deletions of the three ERCEs reveal partial redundancy and interdependency among the sites. RT profiles of two replicates for the indicated deletions are presented with deleted allele in red, and WT allele in black. Profiles from two different clones are presented for all but the bc deletion.

### ERCEs regulates transcriptional activity

Due to the presence of enhancer-like features and binding sites for transcription factors within ERCEs, we evaluated the consequence of deleting these enhancer-like elements on transcription by Bru-seq (Paulsen et al., 2014), which captures nascent RNA transcripts (**Figure 5a** and **Supplemental Figure 7**). Transcription throughout the domain was nearly eliminated from within the abc deleted alleles, and a reduction in gene transcription was observed in other pair-wise deletions (**Figure 5a** and **Supplemental Figure 7**). Since each ERCE contains a TSS, transcription cessation on those genes was expected. However, the DppA4 gene with its TSS is intact but its transcription was eliminated in the abc deletion, suggesting that one or more ERCEs are necessary for expression of the DppA4 gene. We also observed the presence of a new TSS (not previously detected by CAGE) within the Morc1 gene after deletion of its original TSS (**Figure 5a**, bc deletion). Thirdly, we observed decreased transcription of the full-length Morc1 gene in the site c deletion, which deleted an intragenic segment of this gene (**Figure 5a** and **Supplemental Figure 7**), suggesting site c enhances Morc1 gene expression. These findings suggest that ERCEs contain enhancer activity.

To compare the relative effects of ERCE deletion on RT vs. transcription, we plotted the change (relative to WT) in transcription vs. the change in RT for gene segments as indicated in **Figure 5a** (DppA4: 48283735-48294292, DppA2: 48311882-48319513, Morc1: 48511046-48581359, mm10) that were not included in the deletions and therefore in common between the deleted and undeleted 129 alleles in WT cells (**Figure 5b**). Results revealed that gene transcription was nearly eliminated when the shared gene segment replicated late, while the remaining partial transcription changes were proportionally correlated with changes in RT (Spearman’s correlation =0.70, and Pearson’s correlation= 0.82). This suggests that the effects on RT and transcription are intimately related at this locus. This is not true of the changes in transcription and RT genome-wide (Hiratani et al., 2010; Rivera-Mulia et al., 2015), and plotting RT vs. the Bru-seq nascent transcription for all RT-regulated domains (those known to change RT during cell differentiation) revealed a much weaker correlation (Pearson’s correlation = - 0.01; Spearman’s correlation = 0.2) (**Figure 5c**). We observed that Morc1 continues to transcribe through the intragenic deletion of site c, although the right segment collapses into a late-region-like TTR, suggesting that transcription elongation is not sufficient to maintain early replication (**Supplementary Figure 7c**). It will be interesting to further explore the coupling of RT to transcription now that ERCEs have been identified.

## DISCUSSION

Here, we report the identification of a set of discrete *cis*-elements that regulate replication timing and sub-nuclear compartment in a native chromosomal context that we designate as Early Replication Control Elements (ERCEs). Within this replication domain, ERCEs are partially redundant and interdependent and deletion of all three ERCEs is necessary for a complete domain-wide switch of RT from early to late replication. ERCEs are sites of strong non-CTCF-mediated interaction within the domain and they do not align with the most frequently utilized replication origins. ERCEs harbors a collection of active chromatin features resembling promoter/enhancers, and are implicated in gene expression in the domain. The complex interplay among ERCEs accounts for the difficulties experienced in previous efforts to identify such elements in mammalian cells, and the demonstration that large-scale structural and functional features of chromosomes require discrete *cis*-elements paves the way for a systematic approach to dissect the mechanisms regulating temporal and spatial control of DNA replication and their relationship to gene expression.

### ERCEs: *cis*-regulatory elements linking both temporal and spatial control of DNA replication

*Cis*-elements regulating RT in mammalian cells have been sought for many decades. Prior studies involving tandemly integrated BACs or extra-chromosomal vectors provided preliminary evidence for a genetic influence on RT control and sub-nuclear organization (Pope et al., 2012; Sinclair et al., 2010; van de Werken et al., 2017). We recently showed that ~200kb BACs replicate at times similar to their native RT when maintained as extra-chromosomal circles (Sima et al., 2017). While artificial constructs containing combinations of regulatory elements can alter RT in certain chromosomal contexts (Goren et al., 2008; Hassan-Zadeh et al., 2012), it has been challenging to identify functional *cis*-elements *in situ*. Regulatory elements such as late consensus sequences (Yompakdee and Huberman, 2004) or Fkh1,2 binding sites (Knott et al., 2012) have been reported in yeast, no such cis-acting elements have been identified in metazoans. After an extensive catalogue of deletions and inversions, that would have been nearly impossible prior to the advent of CRISPR, we have demonstrated that multiple discrete ERCEs contribute to early replication of a pluripotency-associated domain in mESCs. ERCEs do not coincide with the most efficient replication origins / initiation zones (**Figure 3b**), consistent with prior studies demonstrating that RT is regulated independently of the number and positions of replication origins (Dileep et al., 2015b). Also consistent with the longstanding and evolutionarily conserved correlation between the temporal order of DNA replication and the spatial segregation of active and inactive chromatin inside the nucleus (Solovei et al., 2016), ERCEs are also responsible for sub-nuclear compartmentalization of replication domains, solidifying the link between temporal and spatial aspects of DNA replication in complex genomes and providing discrete elements of regulation with which to pursue mechanism.

### What are the defining features of ERCEs?

ERCEs seem to have two properties that distinguish them from other sites with similar chromatin/epigenetic features: 1) ERCEs form prominent CTCF-independent interactions that may drive the A/B compartmentalization; 2) ERCEs harbor a portfolio of active chromatin features.

#### ERCEs form prominent CTCF-independent interactions that may drive the A/B compartmentalization

ERCEs correspond to sites that form the most robust CTCF-independent interactions within a domain, which are limited almost exclusively to ERCEs. ERCEs also form strong long-range interactions with ERCE-like elements in other domains. These intra-domain and inter-domain interactions mediated by ERCEs suggest a potential ERCE 3D network along the chromosome, which could potentially drive the physical separation of chromatin with ERCE-like properties from chromatin that lacks these ERCE-like properties, and ultimately the formation of the A/B compartments inside the nucleus. Architectural proteins such as CTCF (Nora et al., 2017) and cohesin (Rao et al., 2017; Schwarzer et al., 2017) are mainly responsible for constructing the smaller units (TADs loop domains, or contact domains), but not RT and A/B compartments. Deletion of CTCF-associated boundaries had no effect on DppA2/4 RT, and architectural protein depletion not only failed to cause widespread RT (this report) or A/B compartment changes (Rao et al., 2017; Schwarzer et al., 2017), but also increased the strength of compartmentalization and super-enhancer mediated interactions (Rao et al., 2017). At DppA2/4 domain, ERCE interactions are not restricted to CTCF/cohesin convergent pairs that form “insulated neighborhoods” (Dowen et al., 2014). These findings uncouple TADs from nuclear compartments and support a potential role of ERCEs in driving A/B compartmentalization independent of architectural proteins.

#### ERCEs harbor a portfolio of active chromatin features

ERCEs contains sites with prominent active chromatin features. The chromatin composition of ERCEs in ESCs includes but may not be limited to: DHSs, P300, H3K27ac, H3K4m1, H3K4m3 or Med1, and pluripotency transcription factor binding sites (Oct4, Sox2, Nanog; OSN). Some of these histone marks are hallmarks of super enhancers, stretch of enhancers, strong enhancers and/or promoters (Bogu et al., 2016; Whyte et al., 2013), although these marks are not sufficient to qualify being ERCEs. Site X is decorated with the enhancer histone marks, but lacks a strong presence of P300 and OSN, which are *trans* factors other than CTCF/cohesin now recognized for their necessity to mediate loops/interactions (Fang et al., 2014). Not all P300/OSN sites necessary though. For example, site Y with similar ERCE chromatin features, but dos not form as robust interactions as ERCEs do, does not contribute to early RT. It is likely that an ensemble of different *trans* factors can achieve ERCE activity with additional mechanisms mediating compartment-driving interactions, which can be modulated by various *trans* factors in distinct cell types.

### Redundancy and interdependency among ERCEs

ERCEs operate in a partially redundant and interdependent fashion to give rise to the domain-wide RT pattern of the DppA2/4 domain. Our results reveal that the DppA2/4 domain can be thought of as two sub-domains, of which the left is redundantly controlled by site a and b, while the right is mainly controlled by site c. What contributes to this difference between ERCEs, and the extent to which domains can split control over their RT (and compartment) during normal cellular differentiation is a question for further investigation. Nonetheless, this redundancy offers a plausible explanation for the previous failure to identify such elements at the human β-globin locus, which was extensively studied for decades as a paradigm for transcription and replication control in vertebrates. The Locus Control Region (LCR) was initially hypothesized to be responsible for RT as a shift to late replication and a correlated loss of active chromatin properties was observed in a naturally occurring deletion (Hispanic deletion) encompassing the major DHSs (Forrester et al., 1990; Kalejta et al., 1998). In addition, the LCR also appeared to be sufficient for directing development-specific RT at ectopic sites *in vivo* (Simon et al., 2001). However, targeted deletion of the LCR at the native locus failed to reproduce the late RT phenotype as observed in the Hispanic deletion, despite its effects on chromatin and a significant reduction in transcription rates (Cimbora et al., 2000; Simon et al., 2001). These findings are in line with ERCE behaviors in our results. ERCEs can be sufficient to maintain early replication but not necessary at the endogenous locus. Our results predict that similar studies to those described here will reveal additional ERCEs at the β-globin locus (Hassan-Zadeh et al., 2012). In yeast, mechanisms regulating RT work by attracting or antagonizing the activity of the essential replication initiation kinase Cdc7 (Rivera-Mulia and Gilbert, 2016b). In larger genomes, ERCEs may promote Cdc7 activity throughout the domain, and the interdependency among ERCEs may reflect the synergistic achievement of increasing local concentrations of Cdc7 activity, leading to quantitative differences in replication time.

### Role of ERCE in transcription control

ERCEs overlap with enhancer/promoters within DppA2/4 domain, and can function as enhancers. Since our capture Hi-C data shows that site a, b, and c are in close spatial proximity, DppA4, DppA2, Morc, and Morc1 are likely positioned in a shared transcription factory. It is possible that ERCEs may mediate the positioning of these genes, as well as the distal nectin/pvrl3 gene, into a shared transcription factory the assembly of which is lost upon ERCE deletion. The fact that ERCEs have an effect on compartmentalization also suggests a potential mechanism explaining why distal active genes have been found to share transcription factories (Osborne et al., 2004). Moreover, transcription changes were quantitatively correlated with RT changes, but this correlation is not reflected genome-wide for this category of genes within domains that could change RT during differentiation (**Figure 6c**) (Rivera-Mulia and Gilbert, 2016b)(Hiratani et al., 2010; Rivera-Mulia et al., 2015). In principle, early replication could be related to the rate of transcription initiation (but not elongation), the frequency of interaction of a gene with the transcription machinery or the amount of genomic space being transcribed. However, no single aspect of transcriptional control correlates with RT genome-wide (Rivera-Mulia and Gilbert, 2016b). As discussed below, we favor a co-regulation model in which RT reflects an independent activity of the factors that regulate transcription, which we believe can account for all the published observations.

**Figure 6.**
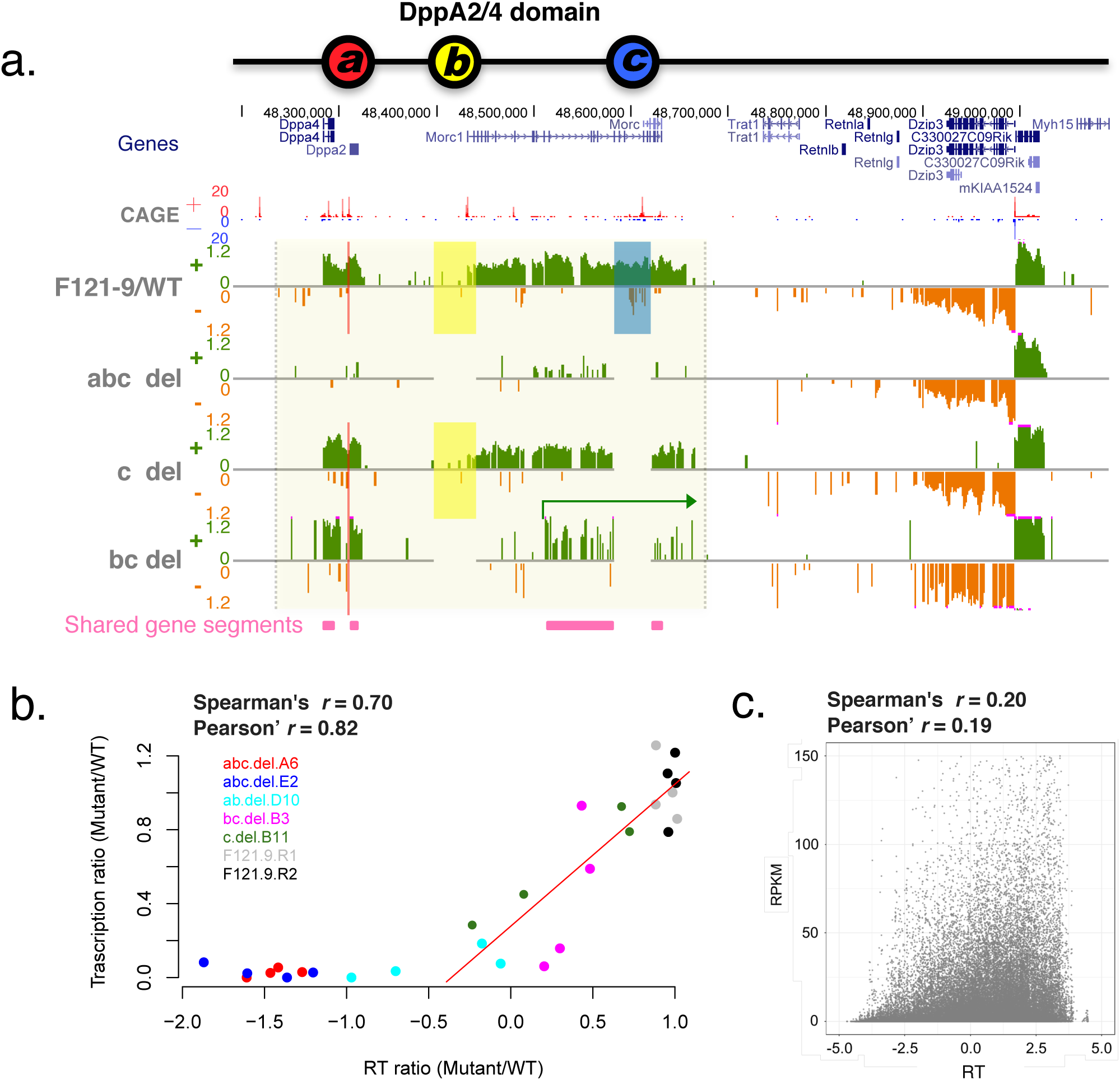
ERCEs are implicated in gene transcription. a. Exemplary Bru-seq profiles for WT and deletions (abc, c and bc deletions). CAGE, Cap Analysis of Gene Expression (ESC). For Bru-seq profiles, negative strand is shown in green, and positive strand in orange. Site a, b, and c are highlighted in red, yellow and blue, respectively. The green arrow indicates transcription of Morc1 gene from a potential alternative TSS upon bc deletion. The pink horizontal bars represent the positions of shared gene segments used for the analysis in Figure 6b. c. Scatter plot of replication timing changes (x-axis) and corresponding transcription changes (y-axis) of shared gene segments within DppA2/4 domain in deletions and WT. A red regression line is added for early replicating segments. d. Scatter plot of the replication timing (x-axis) and transcription levels (y-axis) of all genes from developmental domains.

### A co-regulation model for ERCEs in spatio-temporal regulation of DNA replication and genome function

Causal links between RT, A/B compartmentalization, chromatin composition and transcription have been speculated between these highly correlated events of the genome but not yet been clearly proven. Some evidences suggest that chromatin composition and transcription competency can be directly influenced by RT (Lande-Diner et al., 2009), while other evidence suggests that chromatin composition and transcription can influence RT (Goren et al., 2008). There are also a handful of examples in which these events can be uncoupled (Dileep et al., 2015a). We propose that these highly correlated events not only are regulated by a common set of *cis* elements and *trans* factor, but also involve independent factors contributing to additional layers of complexity for each event. And ERCEs perhaps reflect the root of this co-regulation process, which is at the genetic level. In budding yeast, forkhead transcription factors Fkh1,2 cluster a subset of replication origins to mediate early replication and this activity can be uncoupled from the transcriptional activating function of these TFs using separation of function mutations (Knott et al., 2012; Ostrow et al., 2017). Most likely, cell-type specific TFs are co-opted to promote replication initiation activity within domains whose RT is developmentally regulated in mammalian. Conceivably, clustering of TF-occupied ERCEs may mediate the association of the domain with the A compartment and promotion of Cdc7 activity to increase the probability of replication origin firing within the confines of the topologically-associated replication domain (**Figure 7**). In sum, our findings of the *cis*-elements in RT and A/B compartment control highlight the complexity of *cis* regulation of the linear genome and provide opportunities to further dissect highly related functions from a 3D perspective.

**Figure 7.**
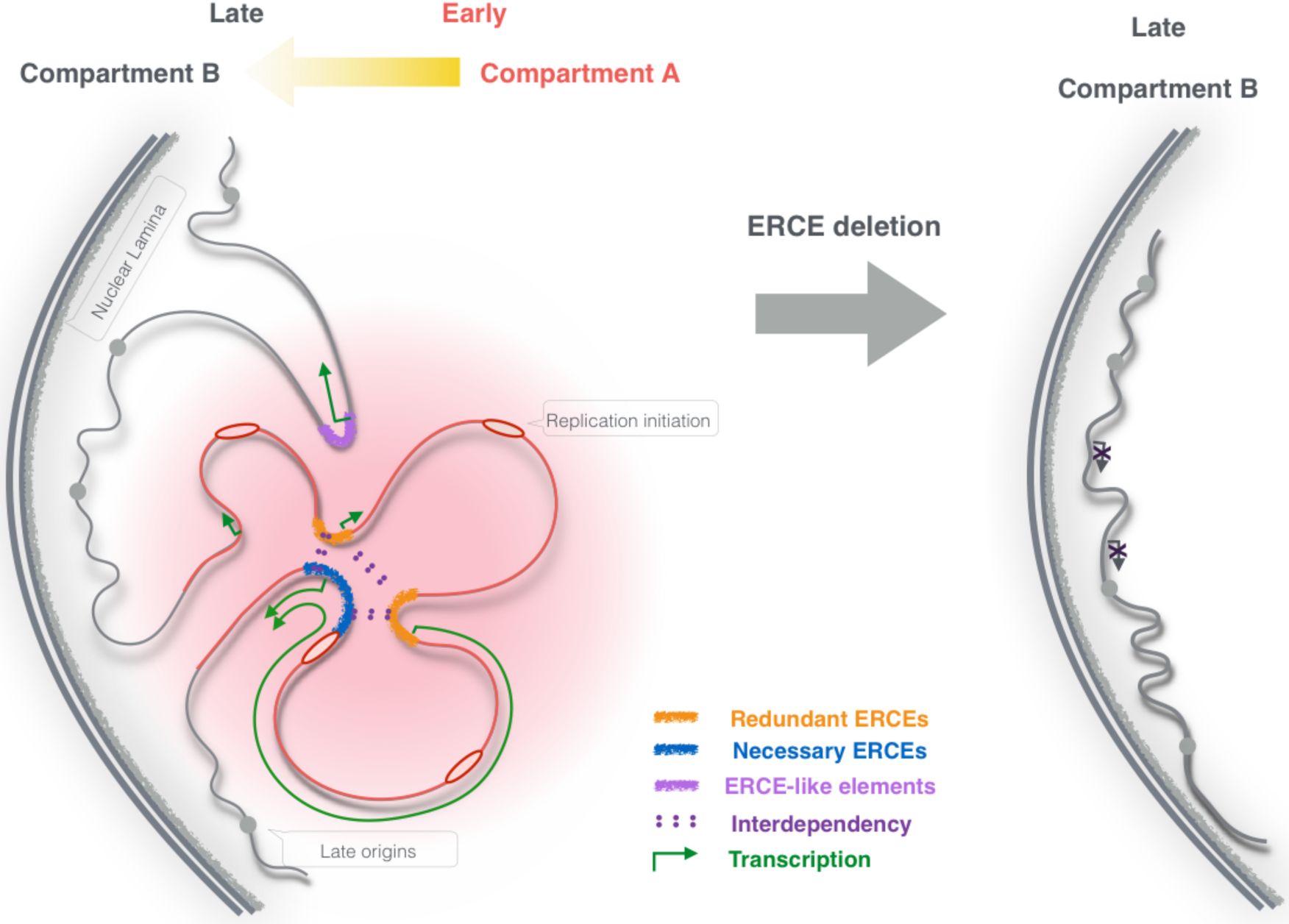
A co-regulation model for ERCEs in spatio-temporal regulation of DNA replication and genome function. A graphic model illustrating ERCEs in regulating RT, A/B compartmentalization and gene transcription. Multiple discrete internal sites (ERCEs) that associated with prominent chromatin features from robust intra-domain interactions with each other and inter-domain interactions with ERCE-like elements, which could drive the formation of compartment A, and lead to early replication. These ERCEs control RT in a partial redundant and inter-dependent manner. Domain boundaries and CTCF do not seem to play an essential role in these processes. ERCEs also activates gene transcription potentially by mediating the assembly of transcription factories. Genetic perturbation of ERCEs leads to loss of compartment A association, late replication, and loss of transcription activity, highlighting *cis* regulation on genome function.

## STAR METHODS

### Cell culture

Mouse ESCs were maintained on 1% gelatin coated dishes. 46C was grown in GMEM supplemented with 10% FBS and LIF; hybrid cells (Cas/129 and V6.5) in DMEM supplemented with 2i plus LIF.

### CRISPR mediated genome editing

CRISPR gRNA (listed in Supplementary Table 2) were designed using http://crispr.mit.edu/, and cloned into pX330. CRISPR mediated genetic engineering pipeline was carried out as previously described with modifications (Byrne et al., 2015). CRISPR plasmids were nucleofected to mESCs Lonza, P3 primary kit) along with a pCAG-mCherry plasmid, and colonies were screened by polymerase chain reaction (PCR) using primers across the breakpoint junctions.

### List of published datasets used in this study

Replication domain boundary calls for DppA2/4 domain were retrieved from (Pope et al., 2014). Hi-C data from ESC and NPC (Bonev et al., 2017) were visualized as heatmaps in HiGlass (Kerpedjiev et al., 2017). Hi-C eigenvector was retrieved from (Nora et al., 2017). Mouse ESC and NPC laminB1 DamID (Peric-Hupkes et al., 2010); nucRNA (Rivera-Mulia et al., 2017), DNaseI HS, H3K27ac, H3K4m1,H3K4m3,H3K36m3, P300 ChIP (Yue et al., 2014), CTCF ChIP (Nora et al., 2017; Yue et al., 2014), SMC1-ChIAPET (Dowen et al., 2014), SMC1-HiChIP (Mumbach et al., 2016), Med1 (Whyte et al., 2013), Oct4, Sox2, Nanog (King and Klose, 2017), CAGE coverage (Arner et al., 2015), SNS mapped origins (Cayrou et al., 2015) used in this study were visualized in UCSC genome browser (Kent et al., 2002).

### Repli-seq

Repli-seq is profiled according to previously described (Marchal et al., 2017). Briefly, cells were pulse labeled with BrdU for 2 hours, and BrdU-substituted DNA from 40k early or late S-phase cells was immune-precipitated by anti-BrdU antibody (Becton Dickinson 347580). Repli-seq libraries were constructed using NEBNext Ultra DNA Library Prep Kit for Illumina (E7370) and sequenced on Illumina HiSeq 2500. For inbred cells, reads were mapped to mm10 reference genome using bowtie2 (Langmead and Salzberg, 2012) with quality score trim at 30. For cas/129 hybrid cells, reads were initially mapped to svim genome with quality score above 10, and SNPs parsed to corresponding allele. Reads overlapping SNPs between the two genomes were filtered and sorted to each allele according to the exact nucleotide at the SNP positions. Reads that do not overlap with SNPs or do not match the exact nucleotide for either genome at the SNP position were discarded. For V6.5 mESCs, reads were mapped to mm10 reference genome and sorted to either allele using same method with SNPs between reference genome and svim. Log2 ratios of early vs late read counts were then calculated in 5kb windows, Loess smoothed, and rescaled to the same interquartile range.

### Capture Hi-C

Capture Hi-C enriching 45488946-50879849 on chromosome 16 was performed as previously described (Dryden et al., 2014). Hi-C libraries from 25 million cells were constructed with 4 base cutter MboI, and target enrichment was performed using Agilent SureSelect RNA oligo system with custom designed capture probes (50bp RNA targeting Mbo1 fragments containing at least 1 SNP). Capture Hi-C libraries were then sequenced on Illumina NextSeq 550 with 150bp pair end. Reads were trimmed by Trim Galore! (https://github.com/FelixKrueger/TrimGalore); mapped using HiCUP v0.5.9 (Wingett et al., 2015), and allele parsed using SNPsplit v0.3.0 (Krueger and Andrews, 2016). Capture Hi-C heatmaps were plotted in R using interaction matrix for 5kb windows. Significant interactions were called using FitHiC (Ay et al., 2014), and visualized as Arcs in WashU browser.

### 4C-seq

4C-seq was performed as previously described (Splinter et al., 2012). 10 million cells were fixed in 1% formaldehyde, and chromatin was digested using HindIII prior to proximity ligation. After reverse-crosslinking, DNA was digested with DpnII prior to circularization. 4C libraries were prepared using inverse-PCR primers containing Illumina adapters that anneal to each bait region (sequences are listed in the supplementary table 4). Libraries were sequenced using 50 bp single-end sequencing on Illumina HiSeq 2500. Reads were mapped to mm10 mouse reference genome and in the case of cas/129 hybrid mESCs, SNP parsed to each allele using same method as described in repli-seq.

### Bru-seq

Bru-seq was performed as previously described (Paulsen et al., 2014). Cells was pulse labeled with 2 mM bromouridine (Bru) for 30 minutes, and nascent RNA was pulled down using anti-BrdU antibodies conjugated magnetic beads. Strand specific cDNA libraries were constructed using 300ng Bru-labeled RNA and sequenced on Illumina HiSeq 2500 with 100bp single end. Reads were mapped using bowtie2, and allele-specific analysis was performed using harp (http://github.com/dvera/harp). RPM-normalized read densities were calculated with bedtools2 (Quinlan and Hall, 2010) genomecov, using a scaling factor of 1000000/(number of parsed reads in library).

## ACCESSION NUMBERS

The Repli-seq, capture Hi-C, 4C-seq, and Bru-seq data from this study have been submitted to the NCBI Gene Expression Omnibus (GEO; http://www.ncbi.nlm.nih.gov/geo/) with accession numbers XXX.

## Author Contributions

J.S and D.M.G designed the study, J.S established the CRISPR-mediated genetic engineering pipeline; K.N.K and J.S designed gRNA and cloned the CRISPR plasmids; J.S, C.T.R and J.C.R performed cell culture; B.K.W and J.S performed PCR screening; J.S performed repli-seq and analyzed the data; V.D, D.B performed 4C-seq and A.C, J.S, V.D analyzed the data; J.S. and M.M performed cHi-C, and J.S, A.C, F.A analyzed the data; M.T.P performed Bru-seq, and J.S, A.C, D.V analyzed the data; A.C and F.A constructed the 3D model; E.P.N provided CTCF degron samples (Nora et al., 2017); K.K provided mESCs engineered at *Epha4* locus (Lupiáñez et al., 2015); J.S and D.M.G wrote the manuscript with input from all authors.

## ACKNOWLEDGMENTS

We thank George Church lab for advices on CRISPRology, Stefan Schoenfelder for technical advice on capture Hi-C; Steven Wingett for helping with HiCCUP mapping, Simon Andrews for designing capture Hi-C probes; Felix Krueger for SNPsplit; Ruth Didier for operating florescent-activated cell sorting; Cheryl Pye for CRISPR plasmids cloning and Kristina Poduch for help with PCR screening; Yanming Yang for supporting NGS sequencing; Carley Huffstetler for modifying the figures; Marcel Mechali for sharing SNS origin data before publication, Peiyao Zhao and Andrew Belmont for helpful discussions. F.A. and A.C. were partially funded by Institute Leadership Funds from La Jolla Institute for Allergy and Immunology and by NIH grant R01MH111267. This work was supported by NIH grant R01GM083337 and U54DK107965 (D.M.G.).

## SUPPLEMENTAL FIGURE LEGENDS

**Supplementary Figure 1.**
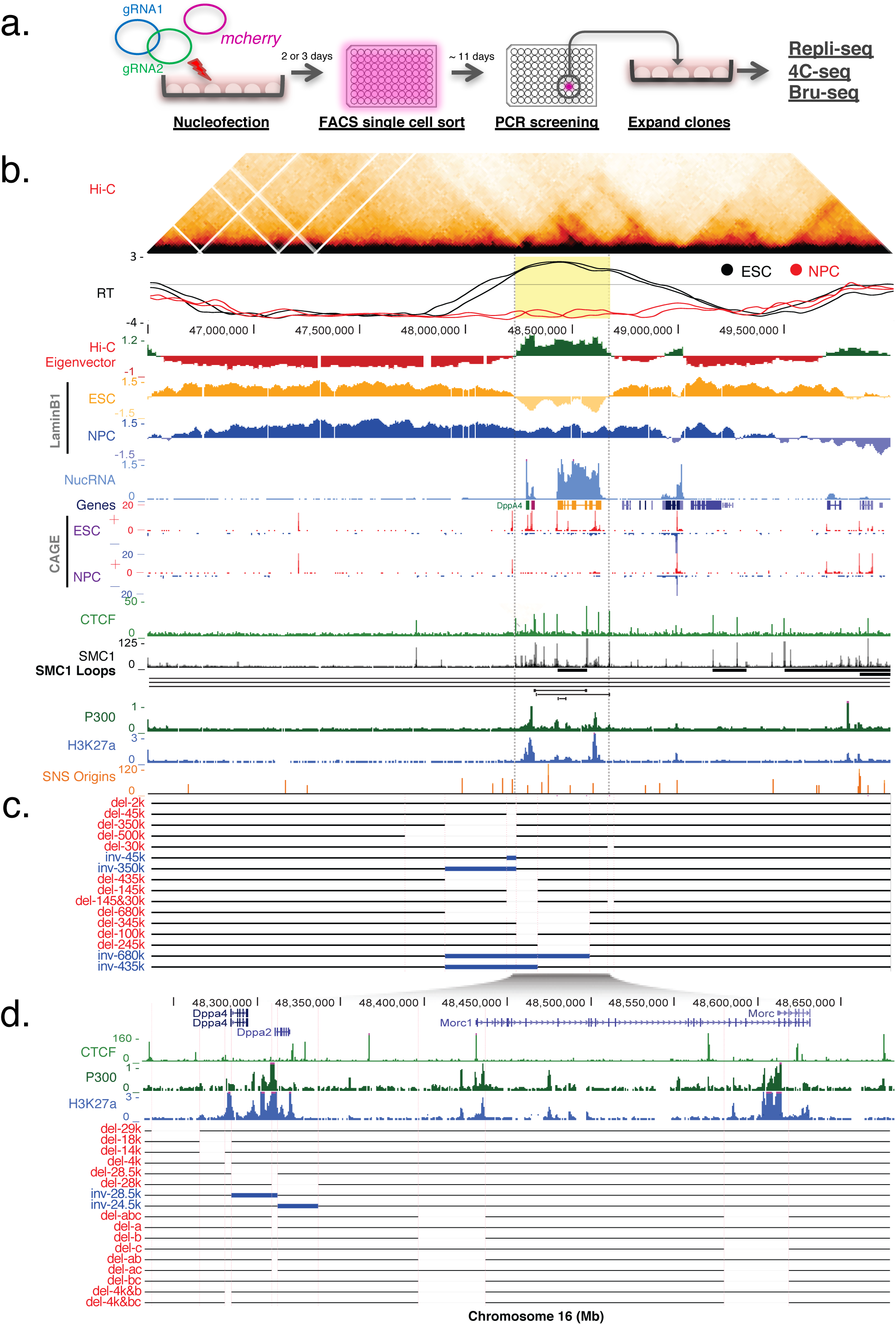
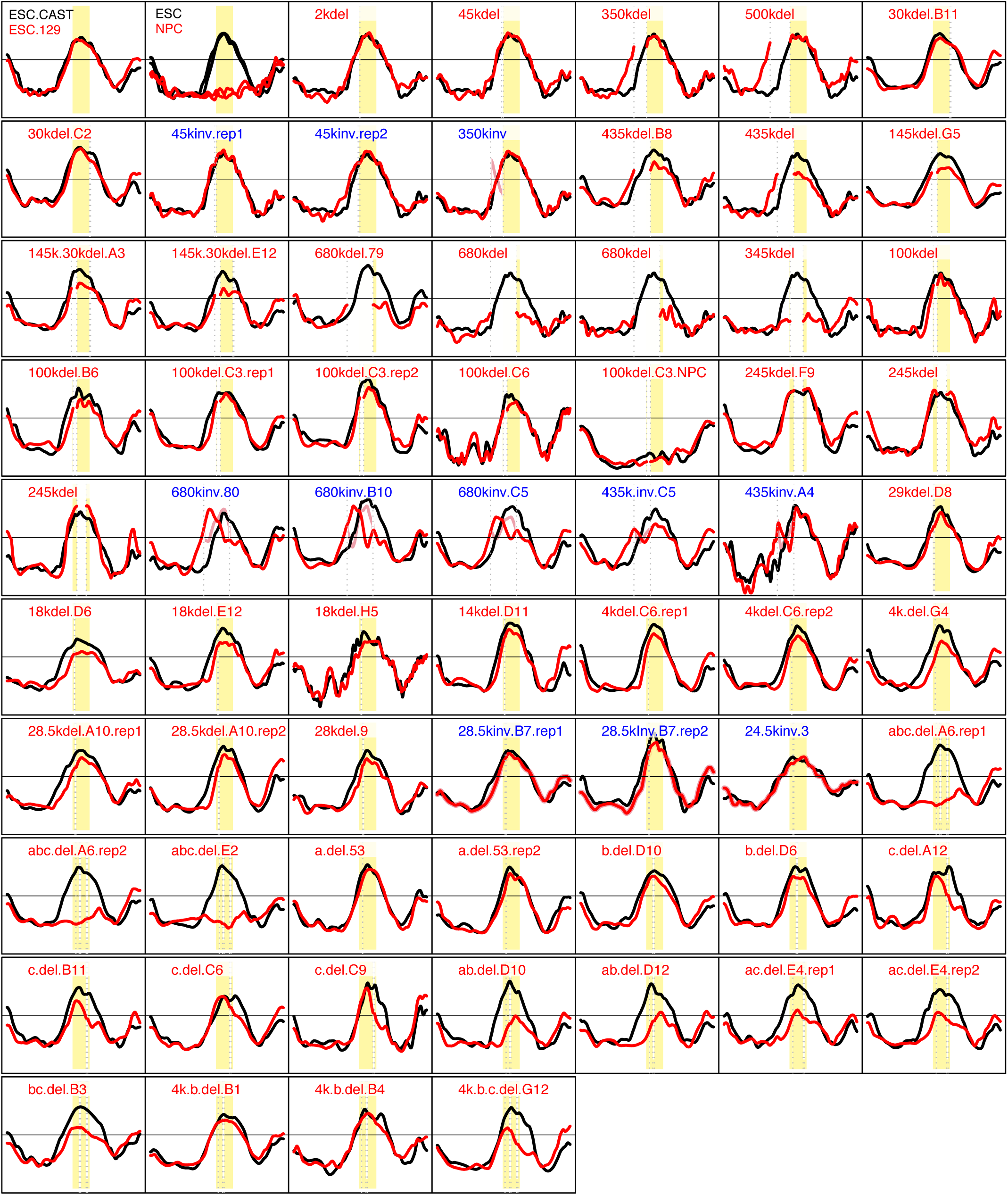
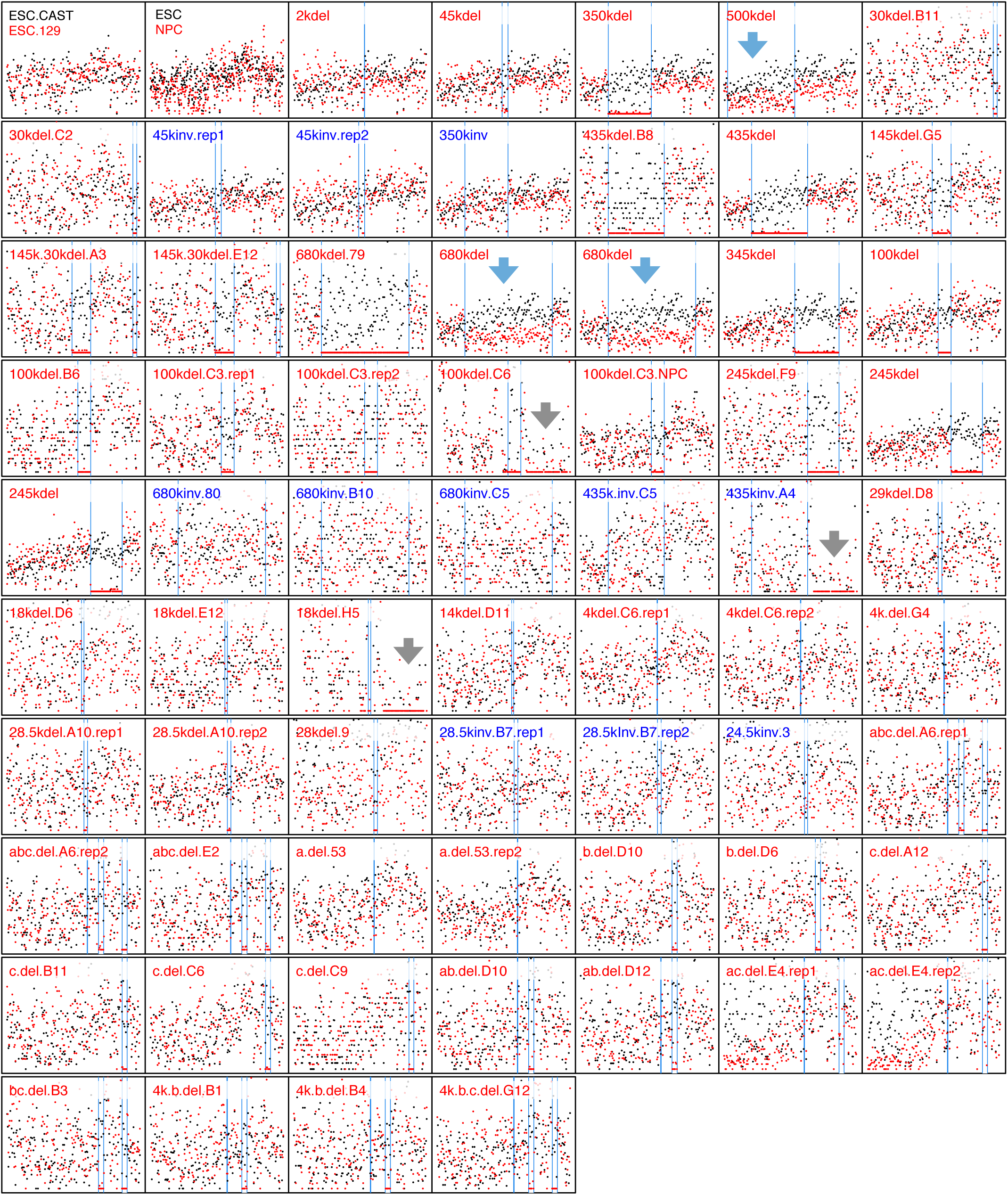
Genetic dissection of the DppA2/4 domain using CRISPRs. a. Experimental pipeline for CRISPR mediated genome engineering in mESCs. Two CRISPR plasmids with guide RNAs targeting each breakpoint were transiently transfected into mESCs along with a mCherry plasmid using nucleofection. Two days post transfection, mCherry positive cells were then sorted as single cells into 96-well plates. Colonies were then screened by PCR using primers spanning the breakpoints and and positive clones were expanded for further analysis (repli-seq, 4C-seq and Bru-seq).
b. DppA2/4 domain chromatin features in ESCs and Neural Precursor Cells (NPCs). Hi-C heatmap is shown for ESCs. RT from both ESC (black) and NPC (red) demonstrate the developmental switch in RT. The DppA2/4 replication domain is highlighted in yellow (chr16:48222368-48668159 from (Pope et al., 2014)), and corresponds well with the DppA2/4 TAD. Hi-C eigenvector is shown at 20kb resolution. Tracks are labeled as follows: LaminB1, LaminB1 DamID; nucRNA, nuclear RNA; CAGE, Cap Analysis of Gene Expression; SMC1 Loops (thick lines), SMC1 ChIA-PET identified loops; SMC1 Loops (thin lines), SMC1 Hi-CHIP identified loops; SNS Origins, short nascent strand mapped replication origins. Data are from ESC unless specified as NPC.
c,d Position of all deletions and inversions generated in this study (see also Supplementary Table 1). c, positions for large size deletions or inversions. d, A zoomed-in view illustrating deletion/Inversion positions of mutants within in the Dppa2/4 domain. The mutants are presented in the order that they are discussed in the main text.
e. RT profiles of all deletions and inversions generated in this study. For mutants (red) generated in inbred mESCs, the 46C WT profile (black) is co-plotted as a control. For mutants generated in hybrid mESCs, the mutant allele is plotted in red, and the WT allele in black, unless otherwise specified. The DppA2/4 domain is shaded in yellow, and deletion positions are indicated by grey dashed lines. The exact genotype configuration of the mutants is listed in Supplementary table 1.
f. Genotype confirmation using repli-seq coverage for all deletions and inversions used in this study. Normalized repli-seq coverage (the sum of RPM from early and late fractions) is plotted in 5kb windows for each cell line. For mutants (red) generated in inbred mESCs, 46C WT (black) is plotted as a control. For mutants (red) generated in hybrid mESCs, the mutant allele is plotted in red, showing a loss in reads for the deleted regions, and the WT allele is plotted in black, unless otherwise specified. Blue vertical lines indicate the breakpoint positions. Heterozygous deletions in 46C inbred ESCs are detected as having one half of the read depth as compared to WT 46C (blue arrows). In V6.5 cells, the region on the right side of DppA2/4 domain is sparse in SNPs between the two alleles, therefore both alleles show 0 coverage in most of the 5kb windows (grey arrows). The exact genotype configuration of the mutants is listed in Supplementary table 1.

**Supplementary Figure 2.**
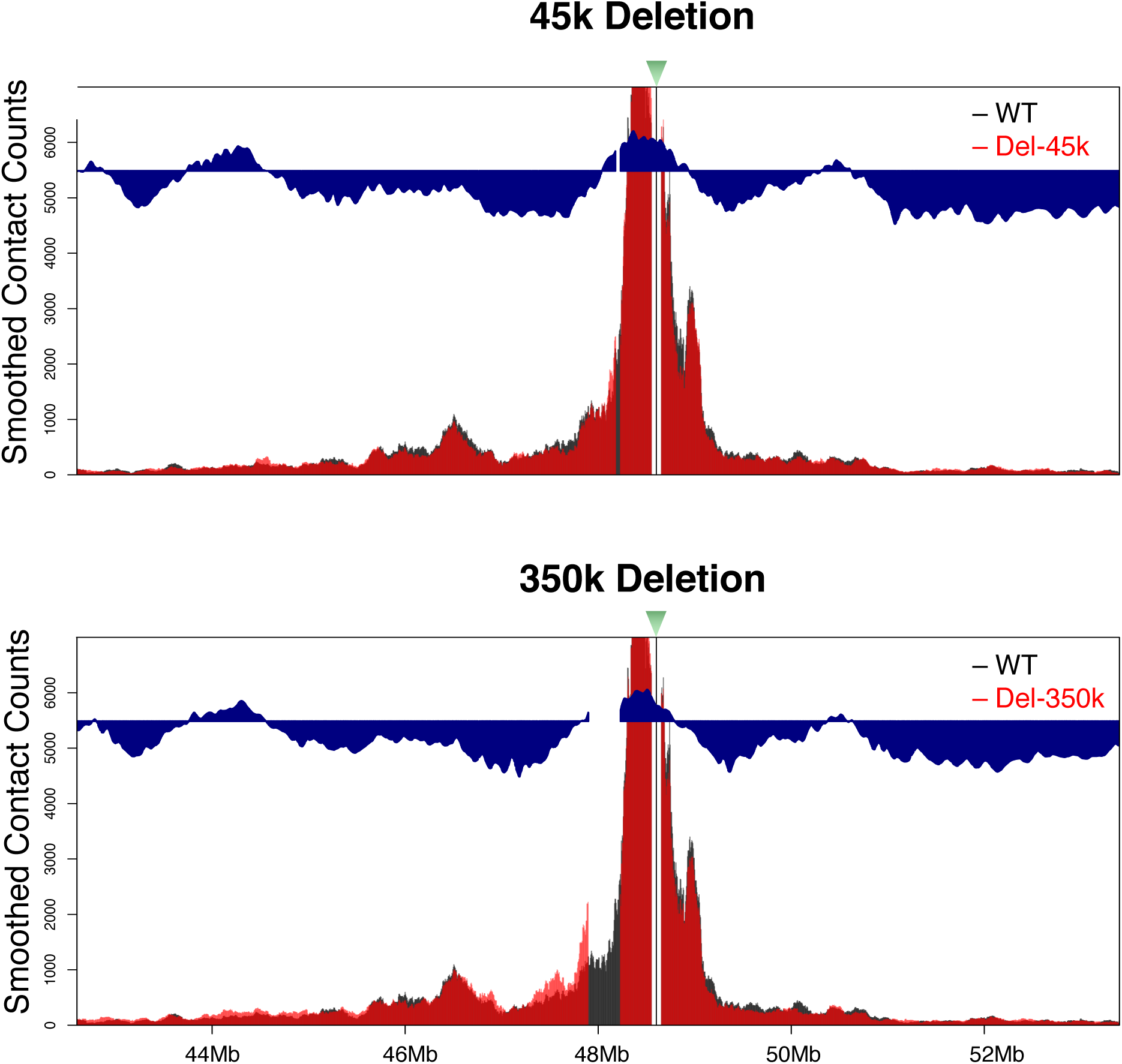
Local 4C interaction pattern of the DppA2/4 domain are not affected after deletion of the CTCF-associated domain boundary. Smoothed 4C contact counts from a bait within DppA2/4 domain (green triangle) are plotted for a 9Mb region in the 45kb (a) or 350kb (b) deletion (red). WT 46C mESCs is plotted in black as a control. Reads within in 50kb of the bait are removed, and the corresponding RT profiles are overlaid as a blue histogram.

**Supplementary Figure 3.**
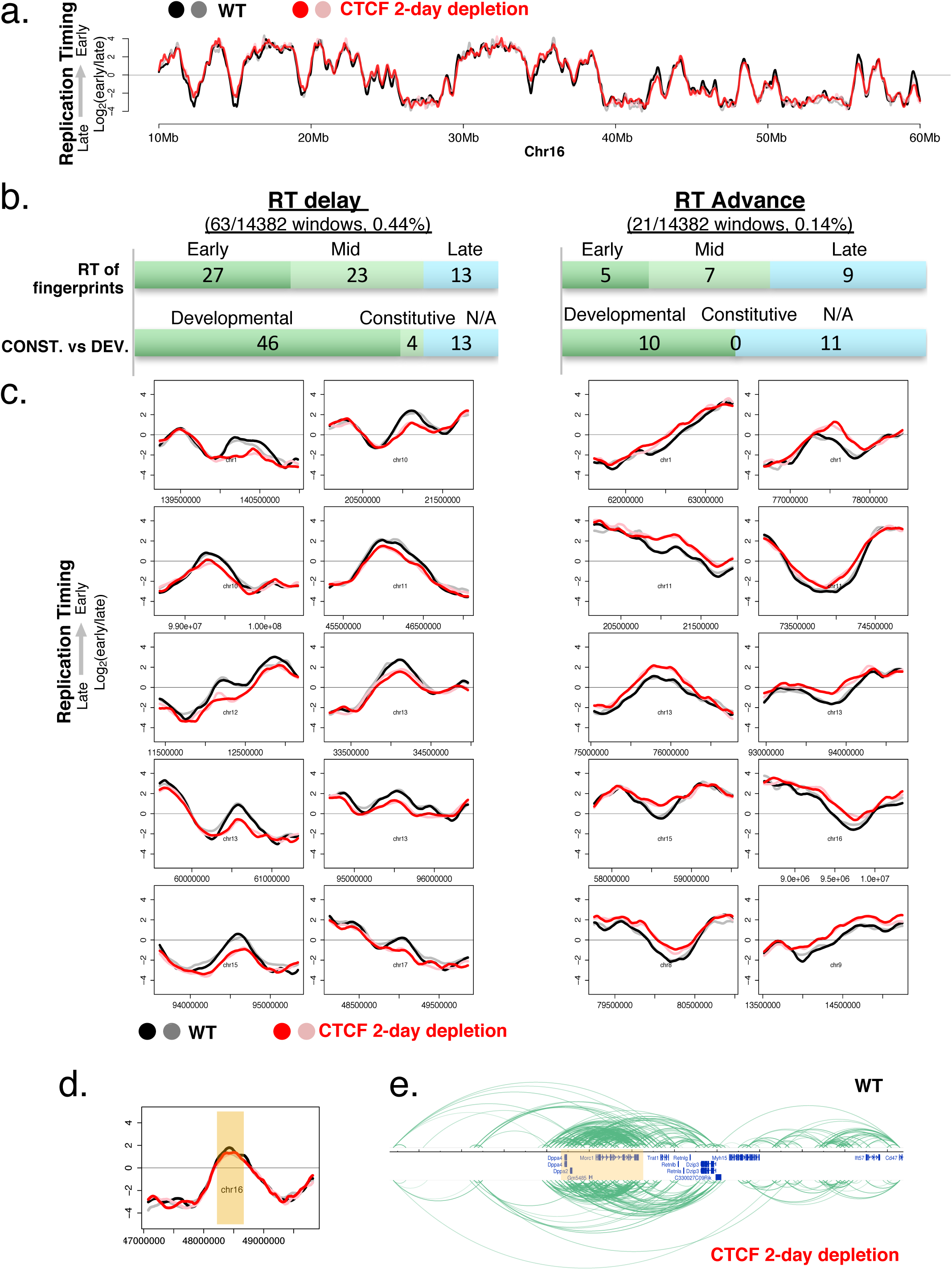
CTCF is not necessary for the genome-wide RT program in mESCs. a. An exemplary RT profile of a 50Mb region of chromosome 16 in WT (black and grey lines), and 2-day CTCF depletion samples (red and pink lines).
b. A summary of CTCF 2-day depletion RT fingerprints in mESCs. For RT values of the RT fingerprint regions, early is defined as RT> 0.5, mid is between −0.5 and 0.5, and late RT<-0.5. “CONST. vs DEV” indicates the number of fingerprints in either constitutive or developmentally regulated domains. Constitutive and developmental domains are defined as in Dileep et al., 2015b. N/A, could not determine.
c. Examples of localized RT changes upon 2-day CTCF depletion.
d. DppA2/4 domain before (black and grey) and after 2 day CTCF depletion (red and pink). DppA2/4 domain is shaded in gold.
e. Arc plot of Hi-C interactions surrounding DppA2/4 domain before (upper panel) and after 2-day CTCF depletion (lower panel). The DppA2/4 domain (highlighted in gold) can still be identified in the 2-day CTCF depletion sample.

**Supplementary Figure 4.**
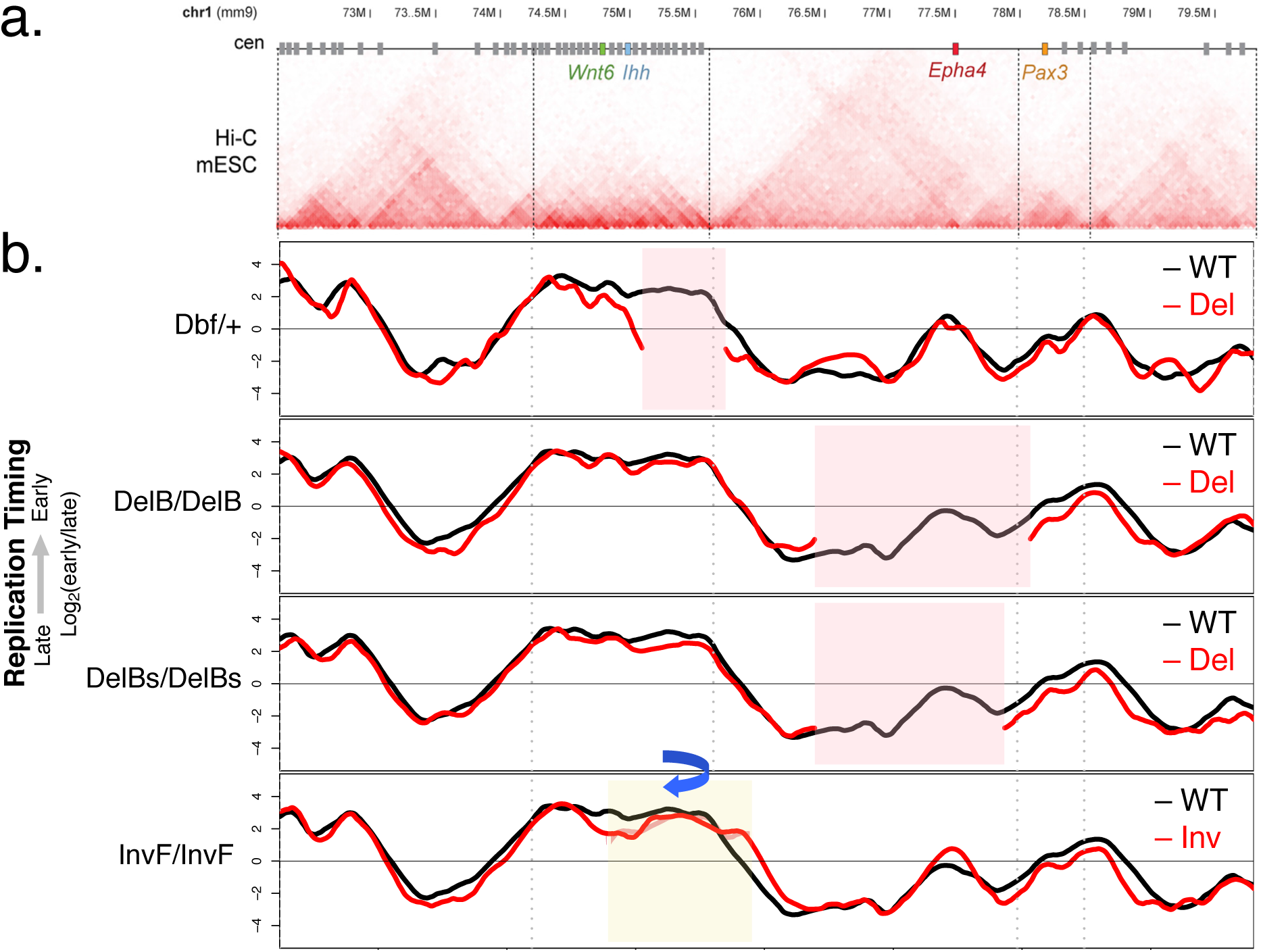
RT of the Wnt6/Ihh and Epha4 domain on chromosome 1 are preserved after deletion or inversion of large segments containing the TAD boundary. a. Hi-C heatmap demonstrating the chromatin structure of the Wnt6/Ihh and Epha4 domains in mESCs (extracted from Figure2 in (Lupiáñez et al., 2015))
b. RT of the deletions/inversions in mESCs. (deletions or inversions in red and WT in black). Deleted regions are highlighted in pink, and inverted region in yellow. RT for inbred Dbf/+ heterozygote was computed *in silico* to simulate the effect on deleted allele (See supplementary methods).

**Supplementary Figure 5.**
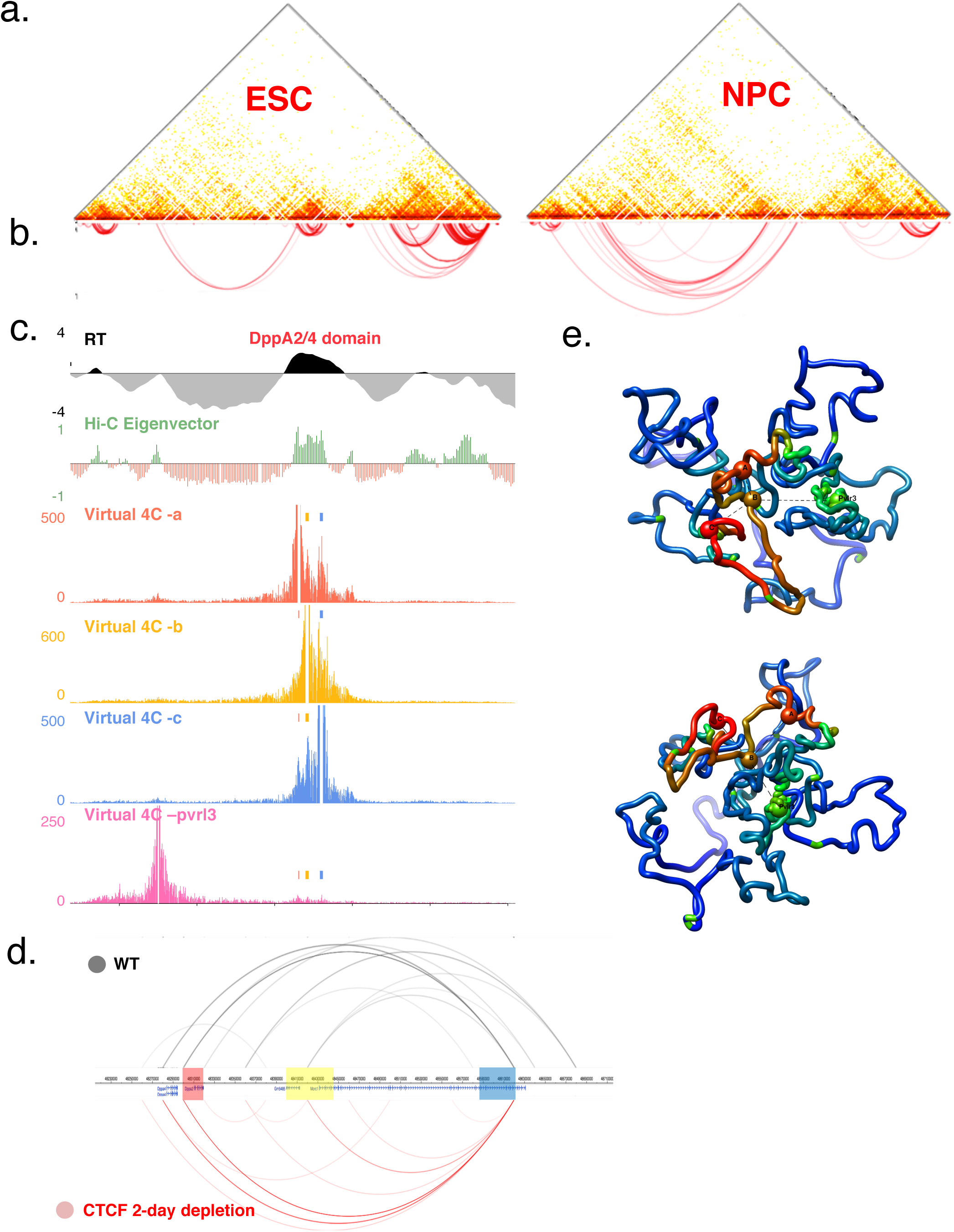
Interactions among site a, b and c are pluripotency specific, CTCF independent, and the most prominent pairs within the DppA2/4 domain. a. Capture Hi-C heatmap of a ~5Mb region containing DppA2/4 domain in ESCs and NPCs, demonstrating the dynamic changes of chromatin interactions during ESC differentiation to NPC.
b. Arc plots of Fit-HiC derived significant interactions in ESC and NPC (FDR < 0.01).
c. Virtual 4C profiles from the viewpoints of either site a, b, c or the pvlr3/nectin3 promoter region at larger Y scale than in figure 4a to demonstrate the interactions among them.
d. Interactions between sites a,b,c are CTCF-independent. Arc plots demonstrate the top 10 Hi-C interaction pairs ranked by FDR for samples before (black) and after Auxin induced CTCF degradation (red). Hi-C data is retrieved from (Nora et al., 2017) with 20kb resolution. Windows containing site a,b,c are highlighted in red, yellow and blue respectively. Interactions between site a, b, and c persist after CTCF depletion.
e. Chrom3D model of the DppA2/4 domain demonstrating spatial proximity between ERCEs and ERCE-like elements. Representation of the Chrom3D model for the DppA2/4 domain 5Mb region from two different angles. Lamin B1 association is reflected as red (lamina depletion) to blue (lamina association) color gradient. Site a,b,c, as well as the immediate ERCE-like element Pvlr3/Nectin3 outside the DppA2/4 domain are indicated as globules, and demonstrate spatial proximity. Grey dashed lines indicate FitHiC significant interactions between the ERCEs and ERCE-like elements.

**Supplementary Figure 6.**
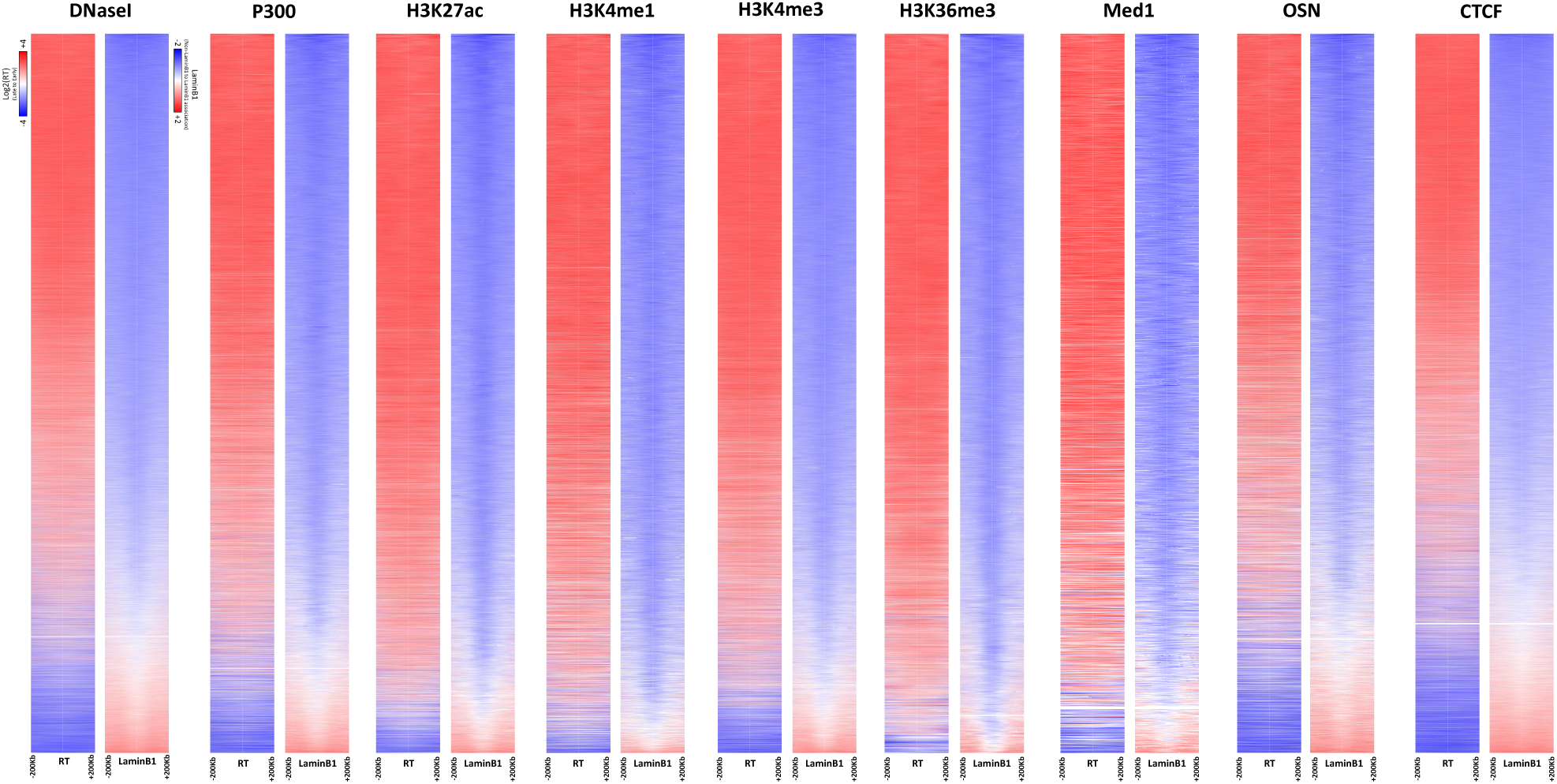
Genome-wide association of chromatin features enriched in ERCEs with early replication and lamina depletion. Heatmaps demonstrating RT or LaminB1 DamID scores for each chromatin feature. Each row represents a peak in corresponding feature, and is ranked by the enrichment of the feature at the peaks.

**Supplementary Figure 7.**
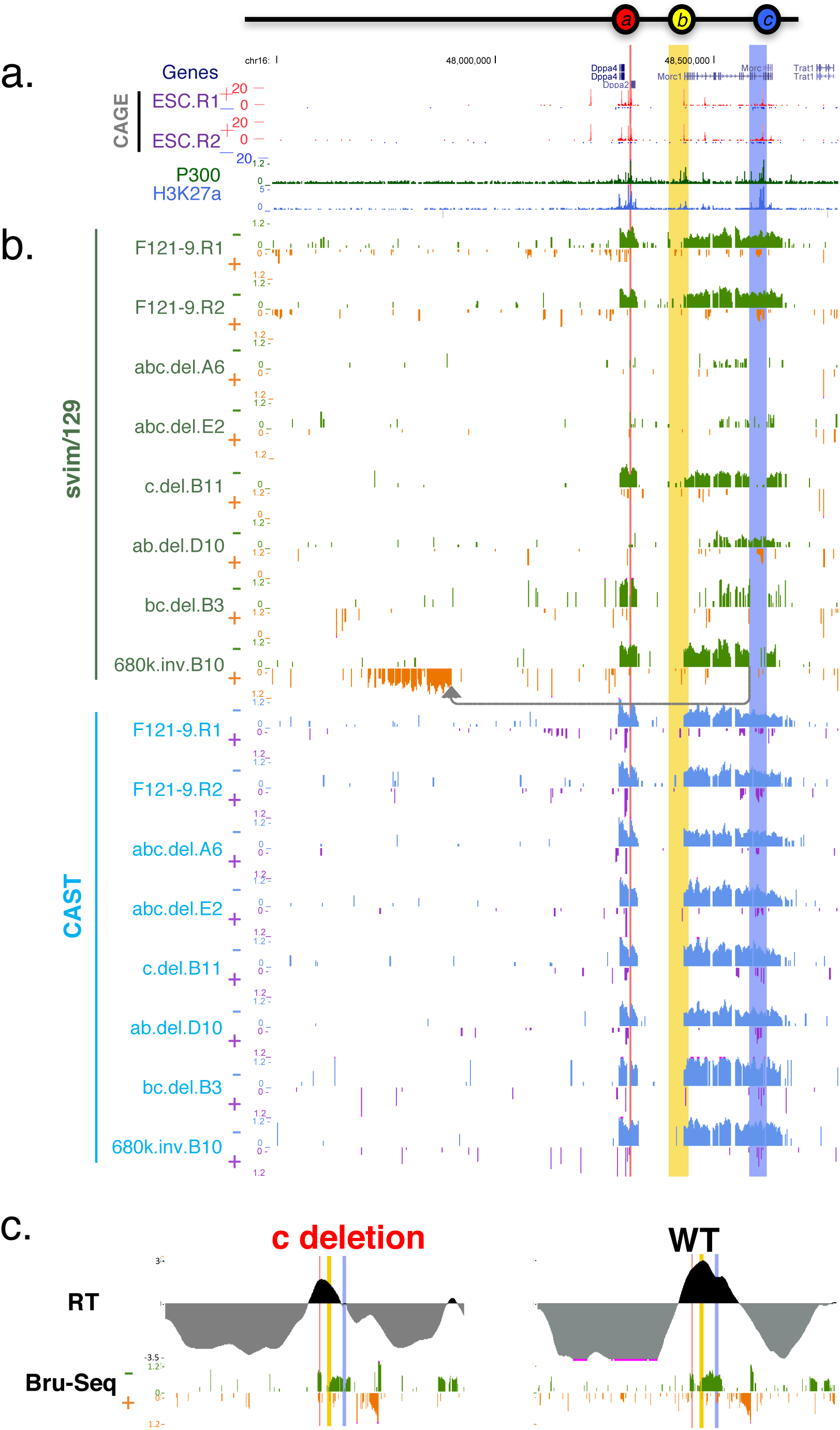
All Bru-seq profiles generated for this study. a. Chromatin features for the plotted region. CAGE, Cap Analysis of Gene Expression. ESC.R1 and ESC.R2 represent two replicates in 46C mESCs.
b. Bru-seq profiles for musculus (svim/129) and the casteneus (CAST) allele of the DppA2/4 locus for WT (R1 and R2 as two replicates), abc deletion (A6 and E2 as two clones), c deletion (clone B11), ab deletion (clone D10), bc deletion (clone B3), and 680k inversion (clone B10). For the 129 allele, the negative strand is shown in green, and the positive strand in orange. For the CAST allele (all WT), the negative strand is plotted in blue, and the positive strand in purple.
c. Left Panel: RT and bru-seq profiles of deletion c demonstrate that Morc1 transcription elongation is not sufficient to maintain early replication, as the gene collapses into a timing transition region (TTR). Right panel: Corresponding profiles are plotted for F121-9 WT.

**Supplementary Table1.** A catalogue of all deletions and inversions generated in this study.

**Supplementary Table2.** A list of CRISPR gRNA primer sequences used in this study.

**Supplementary Table3.** A list of screening primers used in this study.

**Supplementary Table4.** 4C bait sequences used in this study.

## SUPPLEMENTAL EXPERIMENTAL PROCEDURES

### Cell culture

ESC to NPC differentiation was carried out as previously described (Abranches et al., 2009). mESCs were plated on 0.1% gelatin-coated dishes in serum-free ESGRO Complete Clonal Grade medium (Millipore Inc.) at high density. After 24 hours, cells were passaged to RHB-A media (Stem Cell Science Inc.) at the density of 1−10^4^cells/cm^2^ and maintained for 12 days with media changed every other day. CTCF degron mESCs were maintained under the same conditions as in (Nora et al., 2017) and *Epha4* cells as in (Lupiáñez et al., 2015).

### CRISPR mediated genome editing

CRISPR mediated genome editing was performed as previously described (Byrne et al., 2015) with modifications. Two CRISPR plasmids with guide RNAs targeting each breakpoint were transiently transfected into mESCs by nucleofection (Lonza, P3 primary kit) along with a pCAG-mCherry plasmid. At two days post-transfection, mCherry positive cells were then sorted as single cells into 96-well plates. After colonies appear, cells in each well were dissociated and the plate was duplicated; genomic DNA from one of the 96-well plates was prepared as previously described (Ramírez-Solis et al., 1992), and mutations are screened by polymerase chain reaction (PCR) using primers across the breakpoint junctions. PCR fragments were Sanger sequenced to identify allele-specific mutation based on SNPs on both sides of the breakpoints. Positive clones were then further expanded for downstream experiments.

### Repli-seq

Cells were labeled with BrdU (Sigma Aldrich, B5002) for 2 hours at 37°C, fixed in 75% ice-old ethanol with gentle rotation, and sorted into early and late S-phase fractions by flow cytometry based on DNA content. BrdU-substituted DNA from was immune-precipitated using anti-BrdU antibody (Becton Dickinson 347580), and purified after proteinase K digestion overnight. Libraries were prepared using NEBNext Ultra DNA Library Prep Kit for Illumina (E7370), and sequenced on Illumina HiSeq 2500.

### Capture Hi-C

Capture Hi-C was performed as previously described (Dryden et al., 2014). 25 million cells were fixed within 2% formaldehyde at RT fro 10 minutes, and quenched with 0.125M glycine. Cells were then lysed in 50ml ice-cold lysis buffer (10 mM Tris-HCl pH 8, 10 mM NaCl, 0.2% Igepal CA-630, one tablet protease inhibitor cocktail (Roche complete, EDTA-free, 11873580001) on ice for 30 minutes. Collected cell pellet were then incubate in 2.8% SDS with NEBuffer 2 at 37°C for 60 min to remove proteins that were not directly cross-linked to the DNA. SDS was quenched by Triton X-100 at 37°C for 60 min. Chromatin for each sample were then divided into 4 tubes digested with 1500 units of MboI (NEB R0147M) for each tube at 37°C overnight while rotating (950 rpm). To fill in the restriction fragment overhangs and mark the DNA ends with biotin, 6µl 10x NEB2, 2µl H2O, 1.5µl 10 mM dCTP, 1.5µl 10 mM dGTP, 1.5µl 10 mM dTTP, 37.5µl 0.4 mM biotin-14-dATP (Life Technologies 19524-016), and 10μl 5U/μl Klenow (DNA polymerase I large fragment, NEB M0210L) were added to each tube, and incubated for 60 minutes at 37°C. nuclei were then centrifuged down and resuspend in 1ml ligation mix (100µl 10x ligation buffer (NEB B0202S), 10 µl 20mg/ml BSA (NEB B9001S), and 885 µl H2O). For ligation, 5µl 400U/μl T4 DNA ligase (NEB M0202S) was added to each tube for 4 hours at 16°C, followed by 30 min RT. Crosslinks are reversed and protein is degraded by adding 60µl 10 mg/ml proteinase K (Roche 03115879001) per tube and incubating the tubes overnight at 65°C. DNA was then purified by 2 rounds of phenol:chloroform (Sigma P3803) purification, and Hi-C libraries from all 4 tubes for each sample were pooled. PicoGreen (Life Technologies P7589) were used to determine the DNA yields, and Hi-C ligation efficiency was controlled by PCR using cell-type specific primers. Biotin-14-dATP at non-ligated DNA ends is removed from 40μg of Hi-C libraries for each sample with the exonuclease activity of T4 DNA polymerase. Hi-C libraries were divided into 8 tubes, and 5μg of Hi-C library with 0.5μl 10 mg/ml BSA, 5μl 10x NEBuffer 2, 2μl 2.5mM dATP, and 5μl T4 DNA polymerase (NEB M0203L) were added and incubate at 20°C for 4 hours. The reaction is stopped by adding 2 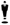 l 0.5 M EDTA pH 8.0 to each tube. DNA were then purified by phenol:chloroform extraction. DNA was then sheared using Covaris E220, end repaired using a home-brew enzyme mix, and size selected between 200 and 650 bp using double-sided SPRI beads. Generate PE adapters were then ligated to the fragments and Biotin-streptavidin pulldown was performed using Dynabeads MyOne Streptavidin C1 beads (Life Technologies 650.01). Hi-C libraries were then amplified using 5-9 PCR cycles using Phusion polymerase (NEB F531). Hi-C library capture was then performed following Agilent SureSelect RNA oligo system protocol, and a final round of PCR amplification (~4 cycles) for capture Hi-C libraries was performed before the libraries were purified using SPRI beads, quantified using KAPA and sequenced with one lane with 150bp pair end on Illumina NextSeq 550.

### 4C-seq

4C-seq was performed as previously described (Splinter et al., 2012) with minor modifications. 10 million cells were fixed in 1% formaldehyde for 10 minutes at room temperature and quenched with glycine (0.125 M final conc.). Nuclei were isolated in lysis buffer (50mM Tris-HCl ph7.5, 150mM NaCl, 5mM EDTA, 0.5% NP-40, 1% TX-100 containing 1X Roche cOmplete Mini protease inhibitors) and dounced 15 times on ice. Nuclei resuspended in 260 µl warm 1X NEB 2.1 buffer were permeablized using 7.5 µl 20% SDS at 65°C for 10 minutes, and then 20 Triton X100 for 60min at 37°C. Chromatin was digested using 500U HindIII (NEB) for 8 hours at 37°C, and the digestion was repeated twice. Enzyme was deactivated at 65°C for 20 mins and digestion as determined by gel electrophoresis. Chromatin samples were diluted and ligated using 8000 CEU NEB T4 DNA Ligase (M0202M) at 16°C overnight. Ligation efficiency was checked by gel electrophoresis. Chromatin was decrosslinked with proteinase K overnight at 65°C, and treated with RNase A at 37°C for 1 hour. DNA was extracted by Phenol:Choroform and precipitated with Ethanol. Purified DNA was treated with 500U NEB DpnII at 37°C for 8 hours, twice. DNA circularization was performed using 8000 CEU NEB T4 DNA Ligase overnight at 16°C. 4C amplification used 3.2 ug 3C template with inverse PCR primers containing Illumina forward and reverse sequencing adapters. PCR was performed using NEB Q5 Hot Start High Fidelity DNA polymerase with the following thermocycler program: 94°C for 2 min; 94°C for 30 sec; 61-65°C for 1 min; 72°C for 3 min; repeat for 29 cycles; 72°C for 5 min; hold at 4°C. 4C-seq libraries were size-selected then quantified using RT-PCR (KAPA Biosystems) and sequenced using 50bp single-end on Illumina HiSeq 2500.

### BrU-seq

Cells were labeled with 2 mM Bromouridine (Aldrich) for 30 min, and lysed in 3ml of Trizol. Total RNA was extracted using chloroform and dissolved in DEPC-treated water. To pull down the Bromouridine incorporated RNA, total RNA was denatured at 80°C for 10 minutes, incubated with anti-BrdU antibodies conjugated magnetic beads (anti-BrdU: BD Biosciences) for 1 hour at room temperature with gently mixing, and washed 3 times with 0.1% BSA in PBS. The beads were resuspended in 35µl DEPC-treated water, mixed and incubated at 96°C for 10 min to elute the Bru-RNA. Strand-specific cDNA libraries were then prepared with the exclusion of polyA RNA isolation. Briefly, 300ng of Bru-RNA was fragmented by mixing with first strand buffer and random primers and incubated at 85°C for 10 min. The first strand cDNA was synthesized in the presence of Actinomycin D to result in strand specific libraries. This cDNA was purified using AMPure RNAclean beads (Beckman Coulter). The second strand cDNA was synthesized and then purified using AMPure XP beads (Beckman Coulter). Next, end repair, adenylation, and adaptor ligation was performed, followed by a size selection in which the cDNA was run on a 3% NuSieve 3:1 agarose gel (Lonza) and gel slices excised in the 300bp region. The gel slices were purified using the QIAEX II Gel Extraction Kit (Qiagen) and then PCR amplified. Resulting libraries were purified with AmPure XP beads, qualitified and sequenced using 100bp single-end on Illumina HiSeq 2500.

### Computation Analysis

#### Determination of CTCF binding site orientation

CTCF MACSs peaks was called with qvalue < 0.05; CTCF MACSs peaks that were present in both datasets (Nora et al., 2017; Yue et al., 2014) were used. CTCF orientation was determined using PWMEnrich package based on its maximum forward or reverse enrichment score. Mouse CTCF motif information was retrieved from http://hocomoco11.autosome.ru/motif/CTCF_MOUSE.H11MO.0.A.

#### Repli-seq coverage calculation for genotype confirmation

Reads from both early and late fractions for a given profile (inbred) or each allele (hybrid cell) are binned at 5kb non-overlapping windows, and normalized to reads per million (RPM).

#### Repli-seq data analysis

For inbred cells, reads were quality trimmed (q score =20) and mapped to mm10 reference genome using bowtie2 (Langmead and Salzberg, 2012). For cas/129 hybrid cells, reads were initially mapped to svim genome with quality score above 10, and SNPs parsed to corresponding allele. Reads overlapping SNPs between the two genomes were filtered and sorted to each allele according to the exact nucleotide at the SNP positions. Reads that do not overlap with SNPs or do not match the exact nucleotide for either genome at the SNP position were discarded. For V6.5 mESCs, reads were mapped to mm10 reference genome and sorted to either allele using same method with SNPs between reference genome and svim. log2 ratios of early vs late read counts were calculated for 5kb non-overlapping windows, Loess smoothed at 400-500kb windows. Coordinates were corrected for deletions by removing the deleted region in Loess smoothing process. Inversions were smoothed according to actual linear distance in the mutants. Chromosome 16 RT profiles for inbred cell lines were quantile normalized. Chromosome 16 RT profiles for hybrid cells were rescaled to the same interquartile range, and mutant allele was compared to WT allele.

#### *in silico* computing of RT for heterozygous mutants in 46C inbred cells

Log2 ratios of heterozygous deletions in inbred mESCs (46C, and the Dbf/+ in *Epha4* deletion series) was calculated as follows: 2 X heterozygous RT – WT.

#### RT Fingerprinting in auxin induced CTCF depletion

RT signatures are identified as previously described at 150kb windows (Ryba et al., 2011). The fingerprints were filtered with qvalue cutoff at 0.01, and RT value difference >1. Constitutive and developmentally regulated domains are defined as in (Dileep et al., 2015).

#### Capture Hi-C data processing

Capture Hi-C reads were trimmed by Trim Galore! (https://github.com/FelixKrueger/TrimGalore); mapped using HiCUP v0.5.9 (Wingett et al., 2015), and allele parsed using SNPsplit v0.3.0 (Krueger and Andrews, 2016) under standard settings. Capture Hi-C interaction matrix was converted in R, and heatmaps were plotted at 5kb windows. Significant interactions were called using FitHiC (Ay et al., 2014), and visualized as Arcs in WashU browser.

#### Genome-wide association of ERCE associated chromatin features to replication timing and laminB1-DamID profiles

Raw sequencing reads for all the individual ChIP-seq datasets from GSE31039 (H3K4me1, H3K4me3, H3K36me3, H3K27me3, and H3K27ac), GSE36027 (P300 and CTCF), and GSE44288 (Oct4, Sox2, Nanog, and Med1) (Whyte et al., 2013) were aligned using bowtie2 program (Langmead and Salzberg, 2012). We allowed two mismatches relative to the reference and only retained the alignments with Phred quality score greater than 30. The datasets were mapped against the mm10 version of the mouse genome. MACS2 (Feng et al., 2012) peak calling was performed using the following settings ‘-q 0.05 –extsize 200 –m 5 50’ against the respective ChIP-seq input files as control. DNaseI peaks were retrieved directly from GSE37074 study and converted to mm10 coordinates from the mm9 assembly using liftOver tool (Tyner et al., 2017). We then mapped the WT lamin B1 (GSE51334) (Pope et al., 2014) and RT signal (this study) surrounding +/-200Kb of each peak from every sample. In case of Oct4, Sox2 and Nanog data; we identified the common triple peak regions among themselves using bedtools (Quinlan and Hall, 2010) and termed them as “OSN” peaks to plot lamin B1 and RT signal.

#### 4C-seq

For inbred cells, reads were quality trimmed (q score =20) and mapped to mm10 reference genome using bowtie2 (Langmead and Salzberg, 2012). For cas/129 hybrid cells, reads were initially mapped to svim genome with quality score above 10, and SNPs parsed to corresponding allele. Reads overlapping SNPs between the two genomes were filtered and sorted to each allele according to the exact nucleotide at the SNP positions. Reads that do not overlap with SNPs or do not match the exact nucleotide for either genome at the SNP position were discarded. 4C reads mapped at HindIII restriction site resolution were further normalized for library sizes across the WT and ABC Del samples for chromosome 16. Library normalization was performed by ‘calcNormFactors’ function from edgeR package that minimizes the log-fold changes between the samples (Robinson and Oshlack, 2010). The scale-factors were calculated using a trimmed mean of M-values (TMM) between each pair of samples. After library normalization, the raw signal from each 4C sample was converted to counts per million (CPM) values for further downstream analysis. The relative enrichment of 4C interactions in a sample was calculated by comparing a sliding window 100 (w=100) restriction sites compared to a background window of 3000 restriction fragments (W=3000). To account false discovery rate (FDR) for each enrichment, random permutation on the dataset was performed to determine the threshold enrichment at which the FDR was 0.01 (Splinter et al., 2012). 4C interactions above the FDR threshold of 0.01 was considered as significant and successively used to measure their RT and eigenvalue feature distribution (excluding the surrounding 48.20-48.65Mb region). For mESC and NPC virtual 4C profile, we first mapped the Hi-C data from (Bonev et al., 2017) at 5Kb resolution on mm10 genome using HiC-Pro pipeline (Servant et al., 2015). Once mapped, we extracted the interaction corresponding to the chr16: 48455,000-48555000 region (equivalent to the +/-50Kb 4C-bait position of 48507782) from both mESC and NPC samples to create the virtual 4C profile. mESC profile was down-sampled to NPC profile for further analysis. We then repeated the same procedure as mentioned for the real 4C data analysis to calculate CPM values and determined the enriched interactions for virtual 4C profiles above the FDR threshold of 0.01 to measure their RT and eigenvalue feature distribution (excluding the surrounding 48.20-48.65Mb region). 4C profile plots were created using custom R scripts. For visualization purpose, we smoothen both the real and virtual 4C profile with a moving average of 300 HindIII site CPM values.

#### BrU-seq

Reads were mapped using bowtie2, and allele-specific analysis was performed using harp (http://github.com/dvera/harp) with default parameters and SNPs for 129/Sv and CAST/Ei from the Mouse Genomes Project, version 5 (Adams et al., 2015). RPM-normalized read densities were calculated with bedtools2 (Quinlan and Hall, 2010) genomecov, using a scaling factor of 1000000/(number of parsed reads in library).

